# A novel Alex3/Gα_q_ protein complex regulating mitochondrial dynamics, dendritic complexity, and neuronal survival

**DOI:** 10.1101/2021.12.09.471902

**Authors:** Ismael Izquierdo-Villalba, Serena Mirra, Yasmina Manso, Antoni Parcerisas, Javier Rubio, Jaume Del Valle, Francisco J. Gil-Bea, Fausto Ulloa, Marina Herrero-Lorenzo, Ester Verdaguer, Cristiane Benincá, Rubén D. Castro-Torres, Elena Rebollo, Gemma Marfany, Carme Auladell, Xavier Navarro, José A. Enríquez, Adolfo López de Munain, Anna M. Aragay, Eduardo Soriano

## Abstract

In neurons, mitochondrial dynamics and trafficking are essential to provide the energy required for neurotransmission and neuronal activity. Recent studies point to GPCR and G proteins as important regulators of mitochondrial dynamics and energy metabolism. Here we show that activation of Gα_q_ negatively regulates mitochondrial dynamics and trafficking in neurons. Gα_q_ interacts with the mitochondrial trafficking protein Alex3. By generating a CNS-specific *armcx3* knock-out mouse line, we demonstrate that Alex3 is required for Gα_q_ effects on mitochondrial dynamics and trafficking, and dendritic growth. *Armcx3*-deficient mice present decreased OXPHOS complex and ER stress response protein levels, which correlate with increased neuronal death, motor neuron and neuromuscular synaptic loss, and severe motor alterations. Finally, we show that Alex3 disassembles from the Miro1/Gα_q_ complex upon calcium rise. These data uncover a novel Alex3/Gα_q_ complex that regulates neuronal mitochondrial dynamics and neuronal death and allows the control of mitochondrial functions by GPCRs.

---

Mitochondria play a key role in producing the energy necessary for cell functioning and survival. Particularly in neurons, mitochondria are essential to provide the high energy supply needed for neurotransmission and neuronal activity. To fulfill this role, neuronal mitochondria are transported along the extensive dendritic and axonal projections through a complex established with the motor proteins kinesin and dynein, the atypical Rho-like GTPases Miro1 and Miro2, and several adaptors including Trak1/2 proteins^1–3^. Calcium-dependent neuronal activity drives disassembly of this complex, causing local mitochondrial arrest at sites of high neuronal activity levels (with a high energy demand)^4–6^. Consistent with this important role in neurons, alterations in mitochondrial trafficking are closely related to neurodegenerative disorders^7–11^.

The *ARMCX3* gene belongs to the *GPRASP* (GPCR-associated sorting protein)/*ARMCX* (armadillo repeat-containing proteins on the X chromosome) family, an Eutherian-exclusive genomic cluster located in chromosome X, which originated by retrotransposition of the ancestor *ARMC10* gene^12–14^. Whereas mutations in members of the *GPRASP* subfamily are associated with delayed pulmonary development and psychiatric and intellectual disabilities^12^, mutations in *ARMCX* subfamily genes are related to tumor progression^12,15^. The *armcx3* and *armc10* genes are highly expressed in neurons, where they localize to mitochondria and regulate mitochondrial trafficking through their interaction with the Miro/Trak complex in a calcium-dependent manner^14,16^. In addition, mitochondrial *armcx1* has been shown to fuel axonal regeneration^17^.

Several studies have shown the presence of G protein-coupled receptors (GPCRs) and G proteins in mitochondria^18,19^. For instance, mitochondrial cannabinoid CB1 receptors control cellular respiration and energy production besides neuronal mitochondrial trafficking and cognition^20–22^. The Gα_q_ protein subfamily, highly expressed in the nervous system, is localized to the outer mitochondria membrane^19,23^. This subfamily mediates the signaling of important metabotropic receptors (e.g., type I glutamatergic and cholinergic muscarinic receptors) and is involved in a number of physiological processes including motor functions, learning, and memory^24–26^. In the mitochondria, Gα_q_ proteins regulate mitochondrial morphology and OXPHOS activity^23^. However, the molecular mechanisms underlying mitochondrial G protein functions remain largely unknown.

## Results

### DREADD-activated Gα_q_ signaling induces neuronal mitochondria arrest

To investigate the role of Gα_q_ in axonal mitochondrial trafficking we used designer receptors exclusively activated by designer drugs (DREADD) as a chemogenetic tool. Hippocampal neurons were co-transfected with mitoGFP and the DsRed-tagged hM3Dq DREADD-Gα_q_ receptor^27,28^. Axonal mitochondria were imaged before and after addition of clozapine N-oxide (CNO) (Fig. 1a, b). Ligand stimulation strongly reduced bi-directional mitochondria motility. A dramatic reduction in the percentage of time in motion in both the anterograde and retrograde directions was observed, besides a marked decrease in the percentage of motile mitochondria and a subtle increase in the number of stops per mitochondrion (Fig. 1b).

**Fig. 1:**
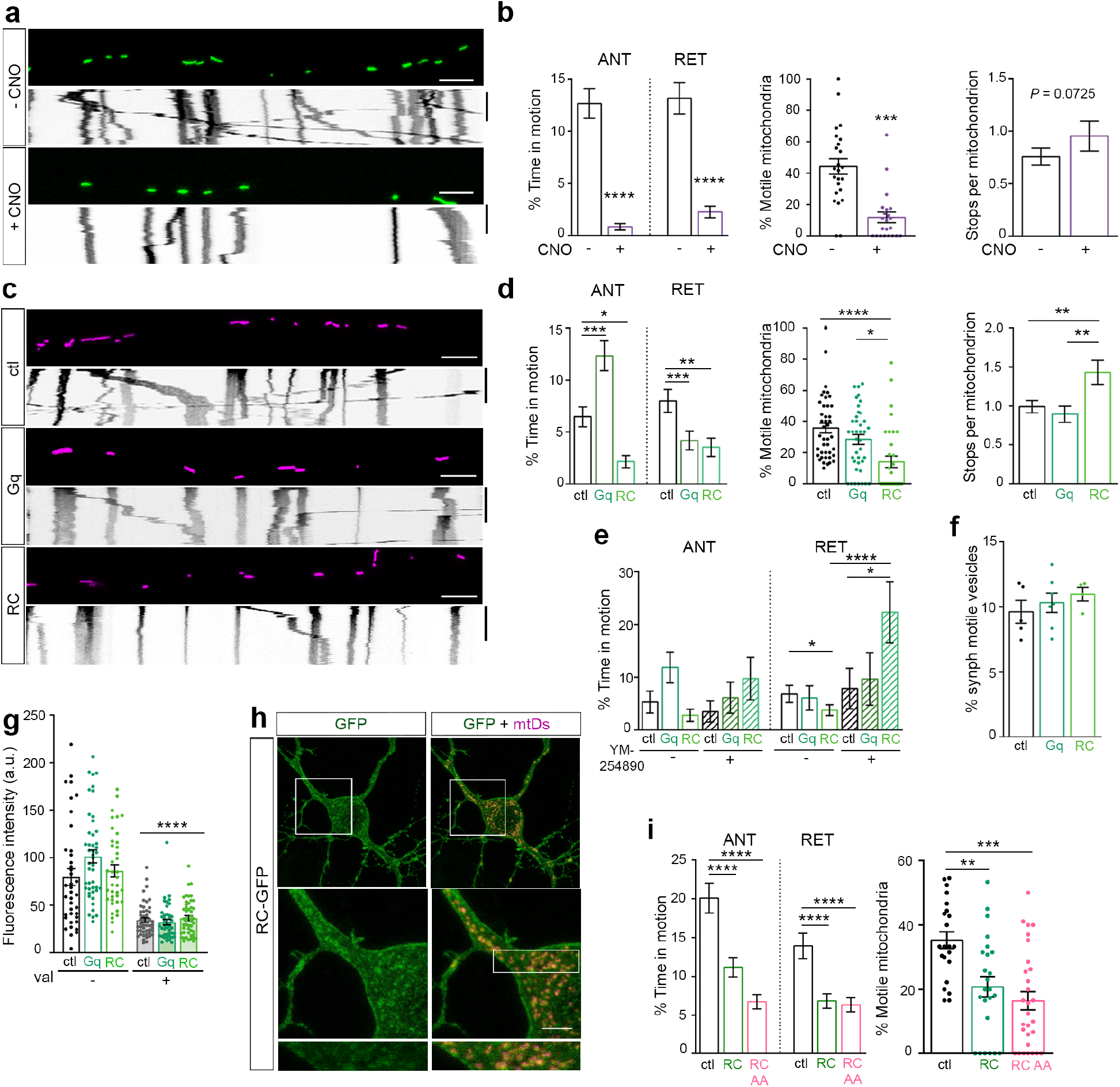
Gα_q_ regulates mitochondria motility in axons. **a**, Representative images and kymographs from axons of live hippocampal neurons expressing the mitochondrial-tagged protein GFP (mitoGFP) and the Gq-specific hM3D DREADD receptor before (-CNO) or 5 min after (+CNO) the addition of 1 μM Clozapine-N-Oxide (CNO), revealing ligand-induced arrest of mitochondria. Images show axonal mitochondria at t = 0 and corresponding kymographs over a 10 min period (y axis). Time bars, 300 s. **b**, Ligand activation (+CNO) reduced “% of time in motion” (TIM) (left) in the anterograde (ANT) and retrograde (RET) directions, decreased “% of motile mitochondria” (MM) (middle), and slightly increased number of stops per mitochondrion (right). n = 387 (-CNO) and 429 (+CNO) total mitochondria from 24 and 22 axons, respectively. **c**, Images and kymographs from axons expressing GFP and mitoDsRed (ctl) with or without Gα_q_ (Gq) or Gα_q_R183C (RC). **d**, % TIM (left), % MM (middle), and number of stops per mitochondrion (right) from (**c**) reflect a decrease in mitochondria motility and more frequent stops upon Gα_q_R183C expression. **a** and **c**, scale bars, 5 μm. Time bars, 300 s. n = 470 (ctl), 390 (Gq), 305 (RC) total mitochondria from 42, 39, and 33 different axons, respectively. **e,** The Gα_q_ inhibitor YM-254890 blocked the effect produced by expression of Gα_q_ and Gα_q_R183C. Neurons transfected as in (**c**) in the presence or absence of 10 μM YM-254890 (INH) or DMSO (ctl). % TIM was calculated from n = 510 (ctl), 452 (Gq), 305 (RC), 29 (ctl+INH), 28 (Gq+INH), and 26 (RC+INH) mitochondria from n = 30, 24, 23, 13, 10, and 20 different axons, respectively. **f,** Quantification of GFP-synaptophysin–tagged synaptic vesicle trafficking in neurons co-expressing mitoDsRed (ctl) with or without Gα_q_ (Gq) or Gα_q_R183C (RC) showed no differences across groups. n = 5 (ctl), 7(Gq), and 4 (RC) different axons. **g,** The fluorescence intensity of MitotrakerGreen from HEK293 cells expressing mitoDsRed (ctl) with or without Gα_q_ (Gq) or Gα_q_R183C (RC) in the presence or absence of 10 μM Val was used as a readout of mitochondrial membrane potential. n =15 (ctl), 17 (Gq), 22 (RC), 27 (ctl+Val), 23 (Gq+Val), and 27 (RC +Val) different cells from three independent experiments. **h,** Images from neurons expressing GFP-Gα_q_R183C and mitoDsRed. Scale bar, 10 μm.**i**, Quantifications of GFP-Gα_q_R183C (RC) and GFP-Gα_q_R183C/R156A/T257A-GFP (RCAA) along with mitoDsRed (ctl) showed similar reductions in % TIM and % MM. n = 336 (ctl), 500 (GFPRC), and 569 (RCAA) mitochondria from n = 22, 24 and 29 different axons, respectively. Data represent mean ± s.e.m. Statistical analyses: **b**, paired non-parametric Wilcoxon test % TIM *P* = 0.0001, % MM *P*= 0.0002; **d, e, f, g** and **i,**Kruskal-Wallis with Dunn correction: **d,**%TIM anterograde (ANT) = 34.54 *P*< 0.0001, retrograde (RET) = 16,87 *P*= 0.0002; %MM = 21.98 *P*< 0.001, number of stops per mitochondrion (Stops) = 11.01 *P*= 0.0041; **e,**%TIM ANT = 10.11 *P*= 0.0722, RET = 23.52 *P*= 0.0003; **f,** Motile vesicles =0.9884 *P*= 0.6327**; g,** FI =68.54 *P*<0.0001, **i,** %TIM ANT = 12.33 *P*= 0.0.0021, RET =6.083 *P*= 0.0478; %MM = 16.90 *P* = 0.0021. \**P*< 0.05, \*\**P*<0.01, \*\*\**P*< 0.001, \*\*\*\**P*<0.0001.

### Gα_q_ regulates mitochondrial trafficking in neurons

The impact of Gα_q_ on mitochondrial motility was analyzed in neurons transfected with cytosolic GFP and bicistronic vectors encoding mitoDsRed and either the constitutively active mutant Gα_q_R183C (RC) or the wild type (Gq) form of Gα_q_. Expression of Gα_q_R183C dramatically reduced anterograde and retrograde mitochondrial trafficking, as well as the percentage of motile mitochondria; in addition, the number of stops per mitochondrion increased, but no alterations in transport velocities were observed (Fig. 1c, d and Supplementary Fig. 1a-c). In contrast, expression of Gα_q_ increased anterograde transport and reduced retrograde trafficking (Fig. 1d).

YM-254890 is a selective inhibitor of Gα_q_ that blocks the exchange of GDP for GTP during Gα_q_ activation^29,30^. Treatment of Gα_q_R183C-transfected neurons with YM-254890 rescued the retrograde transport of mitochondria and reduced the magnitude of defects in anterograde movement observed upon Gα_q_ over-expression (Fig 1e and Supplementary Fig. 1d). Expression of Gα_q_ (either Gα_q_R183C or Gα_q_) had no effect on synaptic vesicle trafficking, as assessed by the expression of GFP-synaptophysin (Fig. 1f and Supplementary Fig. 1e). Further, HEK293 cells transfected with Gα_q_ or Gα_q_R183C and incubated with MitotrakerGreen showed no reduction in fluorescence intensity, but did when treated with Valomycin (Fig. 1g), which suggests that the effects of Gα_q_ are not due to disruption of membrane potential.

The involvement of the canonical Gα_q_ pathway in these effects was evaluated by comparing the GFP-Gα_q_R183C/R156A/T257A^31–33^, a mutant that does not bind or activate PLCβ to GFP-Gα_q_R183C. Interestingly, GFP-Gα_q_R183C/R156A/T257A led to an arrest of mitochondria equivalent to that observed with GFP-Gα_q_R183C (Fig. 1h, i and Supplementary Fig. 1f), but opposite to that observed upon expression of a truncated form of Gα_q_ (GFP-Nt-Gq) (Supplementary Fig. 1g). Together, these findings suggest that Gα_q_ activation induces mitochondrial arrest, while the increase in levels of Gα_q_ enhances anterograde movement, independently of the canonical Gα_q_-PLCβ pathway.

To further analyze the effect of Gα_q_ in mitochondrial trafficking, we knocked down Gα_q_ using specific short hairpin RNAs. Reducing Gα_q_ levels increased retrograde mitochondrial transport and percentage of motile mitochondria, consistently with a decrease in mitochondrial stops (Supplementary Fig. 1h-j), which further supports the involvement of Gα_q_ in regulating mitochondrial motility.

### Gα_q_ interacts specifically with the armadillo domain of Alex3 protein

We next moved on to identify novel binding partners of Gα_q_ that could be involved in mitochondrial dynamics. Endomembranes taken from MEF (WT), NIH3T3 cells, MEF knockout for Gα_q_ and Gα_11_ (KO^23^), and from MEF KO cells expressing Gα_q_ (KO+Gα_q_), were immunoprecipitated and analyzed by mass spectrometry. Among the Gα_q_ proteome, the mitochondrial trafficking protein Alex3 stood out as one of the proteins with the highest number of peptides identified (22 peptides in 3 cell lines) (Supplementary Fig. 2a). Interestingly, Alex3 belongs to the *GPRASP/ARMCX* family, which includes some members identified as putative GPCR interactors^12,13^.

The MS results were then validated in brain homogenates immunoprecipitated with antibodies against either Alex3 or Gα_q_ proteins (Fig. 2a). Endogenous association of Alex3 with Gα_q_ was also confirmed in MEF and SHSY5Y cell extracts (Fig. 2b and Supplementary Fig. 2b). Experiments where Gα_q_ R183C and either myc-Alex3 or GFP-Alex3 were expressed corroborated the interaction between both proteins (Fig. 2c and Supplementary Fig. 2c, d, e). As a control, we immunoprecipitated extracts of cells expressing Flag-Gβ1 and HA-γ2 in the presence of myc-Alex3 (Supplementary Fig. 2f).

**Fig. 2:**
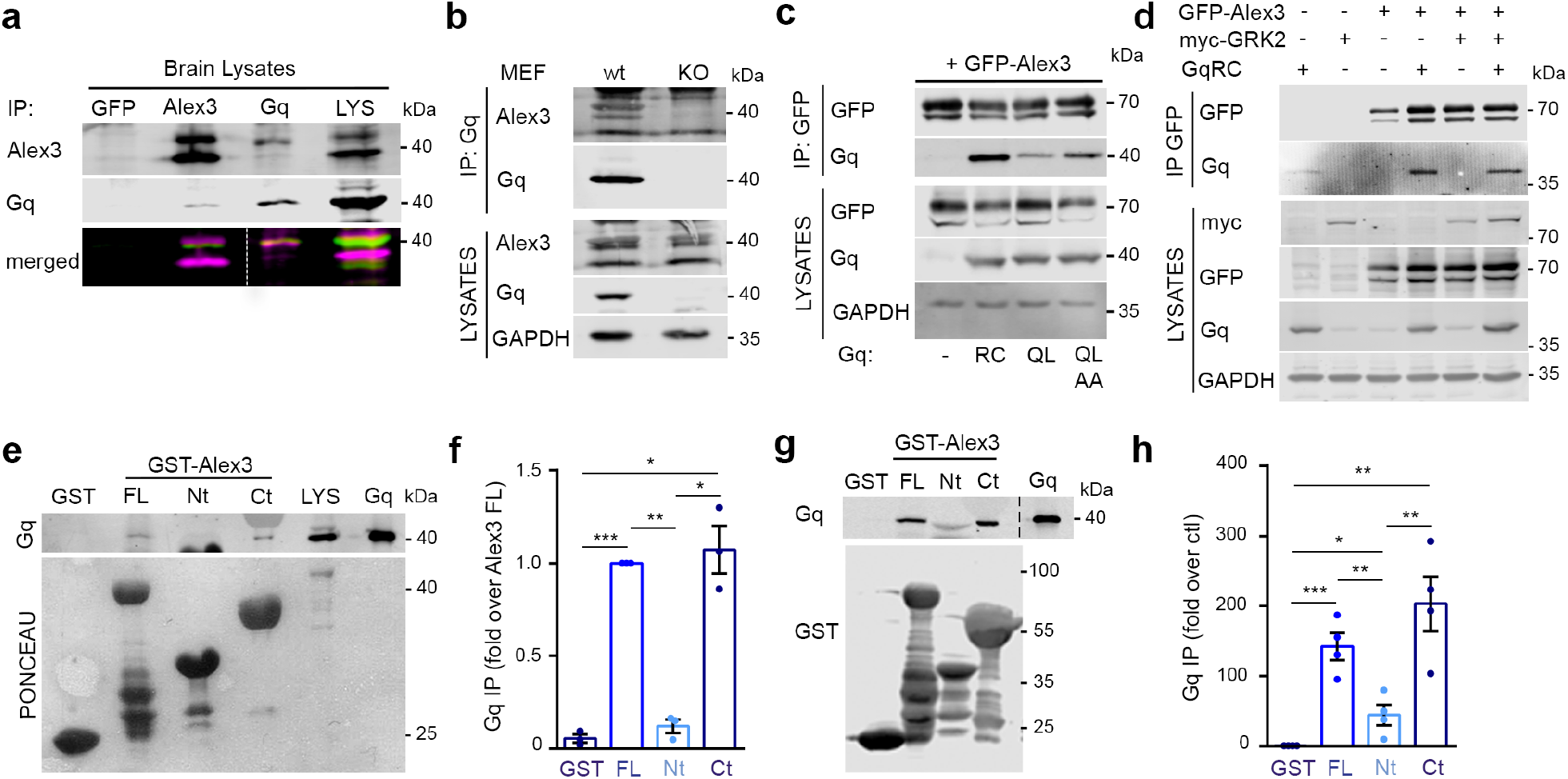
Gα_q_ interacts with Alex3 through its arm domains. **a,** Immunoprecipitation of endogenous proteins from mouse brain extracts (500 μg) using anti-GFP, -Alex3 or -Gα_q_ (E17) antibodies revealed co-precipitation of endogenous Alex3 and Gα_q_ proteins. Homogenates (20 μg) were loaded as input. Alex3 (red) and Gαq (green) are shown. **b,**Endogenous co-immunoprecipitation using anti-Gαq (E17) antibody in MEF WT and MEF Gα_q_/_11_(KO) extracts (1 mg). No immunoprecipitation was detected in control MEF KO cell extracts. **c,** Immunoprecipitation of GFP-Alex3 from HEK293 extracts (700 μg) co-expressing GαqR183C (RC), GαqQ209L (QL), or GαqQ209L/R256A/T257A (QLAA). **d,** GRK2 did not block immunoprecipitation of GFP-Alex3 with Gα_q_R183C from HEK293 extracts (700 μg). **e,** Pull-down of GST-tagged Alex3, Alex3 1-106 (Nt), or Alex3 107-379 (Ct) from SHSY5Y lysates (GST as control). Gq track represents 1 ng of purified Gα_q_ protein, and LYS 20 μg of cell lysates. **f,**Quantification (**e**) represented as fold change relative to Gα_q_ precipitated with the full-length (FL) form. **g,** Pull-down of purified components from 10 ng Gα_q_ with either 10 μg GST or the GST-Alex3 constructs utilized in (**e**). Gq track represents 1 ng of purified Gα_q_ protein. **h**, Quantification from (**g**) represented as fold change relative to Gq in the control condition. Data represent mean ± s.e.m. Statistical analyses: **f** and **h**, one-way ANOVA with Bonferroni correction: **f,**F(38) = 66.82 *P*= 0.0001 n = 3; **h**F(3,12) = 15,81 *P*= 0.0002 n = 4. \**P*< 0.05, \*\**P*< 0.01, \*\*\**P*< 0.001, \*\*\*\**P*< 0.0001.

Alex3 showed reduced or no immunoprecipitation with the constitutive active Gα_q_ Q209L mutant, in contrast to Gα_q_ R183C (Fig. 2c). The difference between these mutants is that Gα_q_ R183C has reduced k(cat) of GTP hydrolysis and still depends on GEF for its activity. Moreover, Alex3 immunoprecipitated with Gα_q_Q209LR183C/R256A/T257A, a mutant that cannot interact with the canonical effectors (Fig. 2c)^31,33^. Expression of GRK2 (a GPCR-specific kinase that binds to Gα_q_ at the canonical effector domain) did not diminish Gα_q_ interaction with Alex3 (Fig. 2d)^34^. These results suggest that Alex3 preferentially binds to the non-active or transitional state of Gα_q_ in a way independent of canonical pathways.

When determining the region of interaction, endogenous Gα_q_ was pulled down with the full-length or the C-terminal (Ct), armadillo-containing region, of Alex3, but weaker or no signal was observed with the N-terminal region from SHSY5Y cell extracts (Fig 2e, f). The region was further delineated with two additional GST-tagged constructs corresponding to the proximal and distal armadillo regions (Supplementary Fig. 2g, h). None of them reached the amount of signal observed with the armadillo domain (Ct). Immunoprecipitation with different GFP-tagged truncated Alex3 proteins^35^ supported these results (Supplementary Fig. 2i). Finally, we took advantage of a purified form of Gα_q_ protein ^36^ to probe the direct interaction with GST-Alex3 full-length and C-terminus in the presence of GDP. Overall, these results confirmed a direct interaction between Gα_q_ and the armadillo domain of Alex3 (Fig. 2g, h).

### CNS-specific inactivation of *armcx3* leads to motor deficits and to abnormal mitochondrial trafficking

To investigate the contribution of Alex3 in Gα_q_-mediated mitochondrial trafficking regulation, we generated a central nervous system (CNS)-specific *armcx3* KO mouse line *(Nestin^Cre^/farmcx3)* (Supplementary Fig. 3a). Western blots of CNS extracts from E14 and P5-7 mouse samples confirmed the absence of Alex3 in *Nestin^Cre^/farmcx3* mice (Fig. 3a-f). Mutant mice were smaller than littermate controls, exhibited severe motor alterations (lack of balance, tremors, and impaired locomotion and coordination) (Supplementary Fig. 3b, c) and usually died around P7-9. We performed further motor behavior-related tests. *Nestin^Cre^/farmcx3* mice presented persistent kyphosis, which might indicate loss of muscle tone, and exhibited deficits in the hind limb suspension and the hind limb clasping tests, which suggests decreased muscle strength (Fig. 3g-i).

**Fig. 3:**
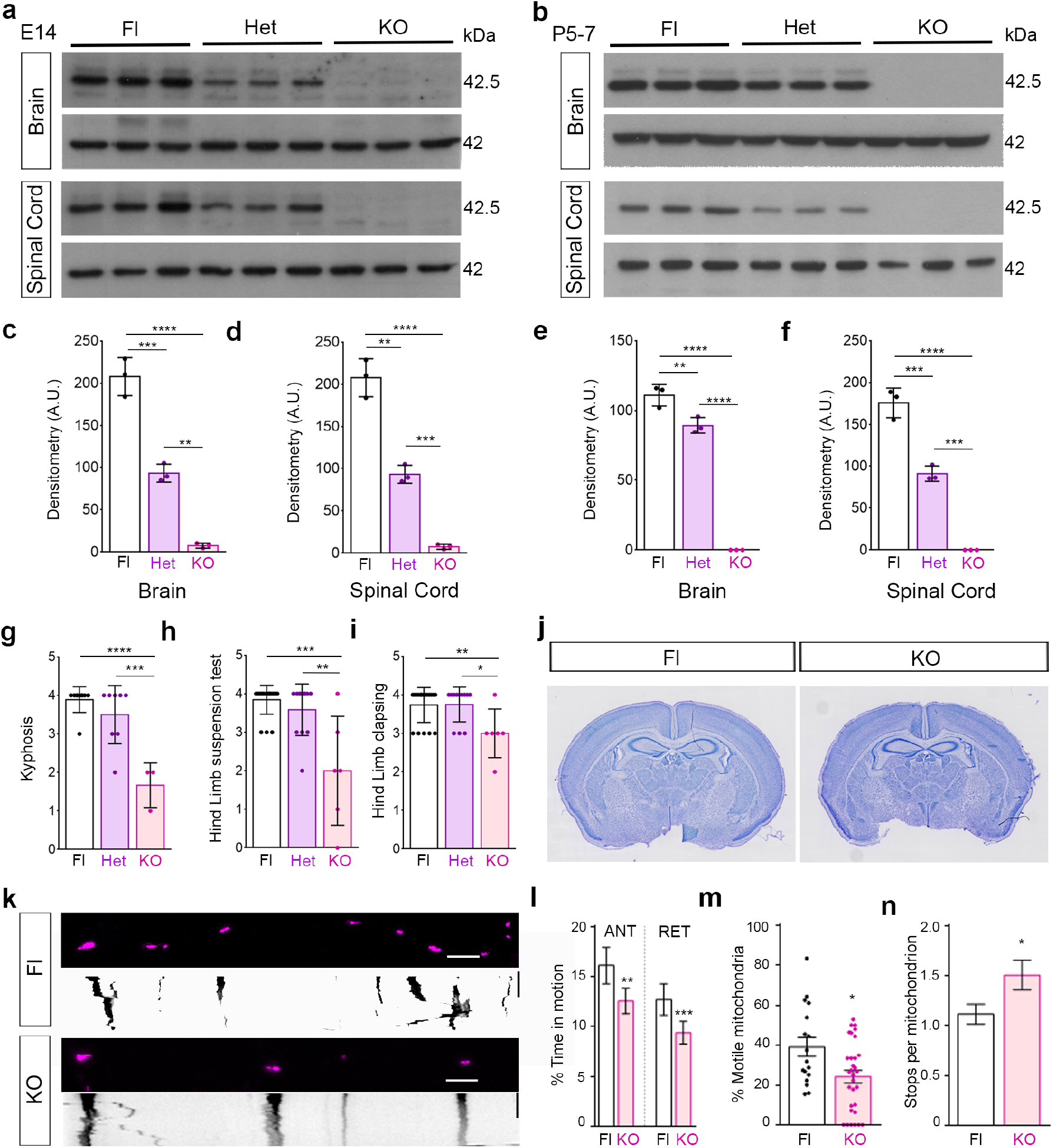
Characterization of the *armcx3* KO mouse line. **a-b**, Representative western blot images of Alex3 in brain and spinal cord from control (Fl), Het, and KO mice at E14 (**a**) and P5-7 (**b**). Actin is used as loading control. **c-f**, Western blot quantification at E14 (**c-d**) and P5-7 (**e-f**). **g-i**, Motor behavior of control (Fl), Het, and KO mice at P5-7; KO mice displayed a kyphosis phenotype, with lower scores (from 4 to 1) in comparison with Fl and Het mice (**g**). KO mice showed lower scores (from 4 to 1) in both hind limb suspension (**h**) and clasping tests (**i**) than did Het and Fl mice. **j**, Representative images of Nissl-stained brain sections from Fl and KO mice. **k**, Images and kymographs from axons of Fl or KO mice neurons expressing mitoDsRed. **l-n**, Graphical representation of “% of time in motion” per mitochondrion (TIM) **(l)**, “% of motile mitochondria” (MM)(**m**), and number of stops per mitochondrion (**n**). Scale bars, 10 μm. Data represent mean ± s.e.m. Statistical analyses: one-way ANOVA with Bonferroni post hoc test: **c,**F(2,6) = 144.9 *P*= 0.0001; **d**, F(2,6) = 119.2 *P*= 0.0001; **e,**F(2, 6) = 331.6 *P*= 0.0001; **f**, F(2, 6) = 174.9 *P*= 0.0001; n = 3 mice per genotype; **g,**F(2, 17) = 17.20 *P*= 0.0001; n = 3-9 mice per genotype; **h**, F(2,34) = 5.932 *P*= 0.0062; n = 6-19 mice per genotype; **i,** Kruskal-Wallis with Dunn = 12.76 *P*= 0.0017; n = 6-19 mice per genotype; **l-n**, U-Mann-Whitney test: %TIM ANT *P*= 0.0087, RET *P*= 0.0007, %MM *P*= 0.0330, Stops *P*= 0.0453. \**P*<0.05, \*\**P*<0.01, \*\*\**P*<0.001, \*\*\*\**P*< 0.0001.

*armcx3* KO mice had slightly smaller brains than littermate controls, which was also noticeable in forebrain histological sections (Supplementary Fig. 3d-n). Despite such a reduction, the overall brain anatomy and organization, including layer distribution in structures such as the cerebral cortex and cerebellum, were similar in *Nestin^Cre^/farmcx3* and control mice (Fig. 3j and Supplementary Fig. 3e-g). We examined the cerebellum of *Nestin^Cre^/farmcx3* mice with anti-Calbindin antibodies, which allow analysis of Purkinje cells at a cellular resolution. Density, body size, and dendritic tree width of Purkinje cells were all reduced in *armcx3* KO mutant mice (Supplementary Fig. 3o-r).

Knock-down of Alex3 has been described to impair mitochondrial trafficking^14^. We thus evaluated the effect of *armcx3* gene deletion on mitochondrial trafficking. Analysis of kymographs of cultured neurons from *Nestin^Cre^/farmcx3* mice showed significant decreases in the percentage of motile mitochondria and time in motion, and increased number of mitochondrial stops, compared with neurons from control mice (Fig. 3k-n). Taken together, our results show that *Nestin^Cre^/farmcx3* mice display severe motor deficits and smaller brains, and confirm previous findings of reduced neuronal mitochondrial trafficking upon Alex3 knock-down^14^.

### Gα_q_ effects on mitochondrial trafficking depend on Alex3

To resolve the contribution of Alex3 in the regulation of mitochondrial trafficking by Gα_q_, we expressed Gα_q_R183C in *Nestin^Cre^/farmcx3* hippocampal neurons. Not only did Gα_q_R183C produce the expected decrease in mitochondria trafficking, but *armcx3* deletion also led to a decrease in motility (Fig. 4a, b). The effects of Gα_q_R183C and *armcx3* deletion were not additive, suggesting that both proteins may be functionally related. Moreover, expression of active Gα_q_ in a *Nestin^Cre^/farmcx3* background rescued both the anterograde movement and the percentage of motile mitochondria (Fig. 4a, b). These results indicate the need of Alex3 to sustain mitochondrial arrest by Gα_q_ in the anterograde transport and suggest a functional link between both proteins.

**Fig. 4:**
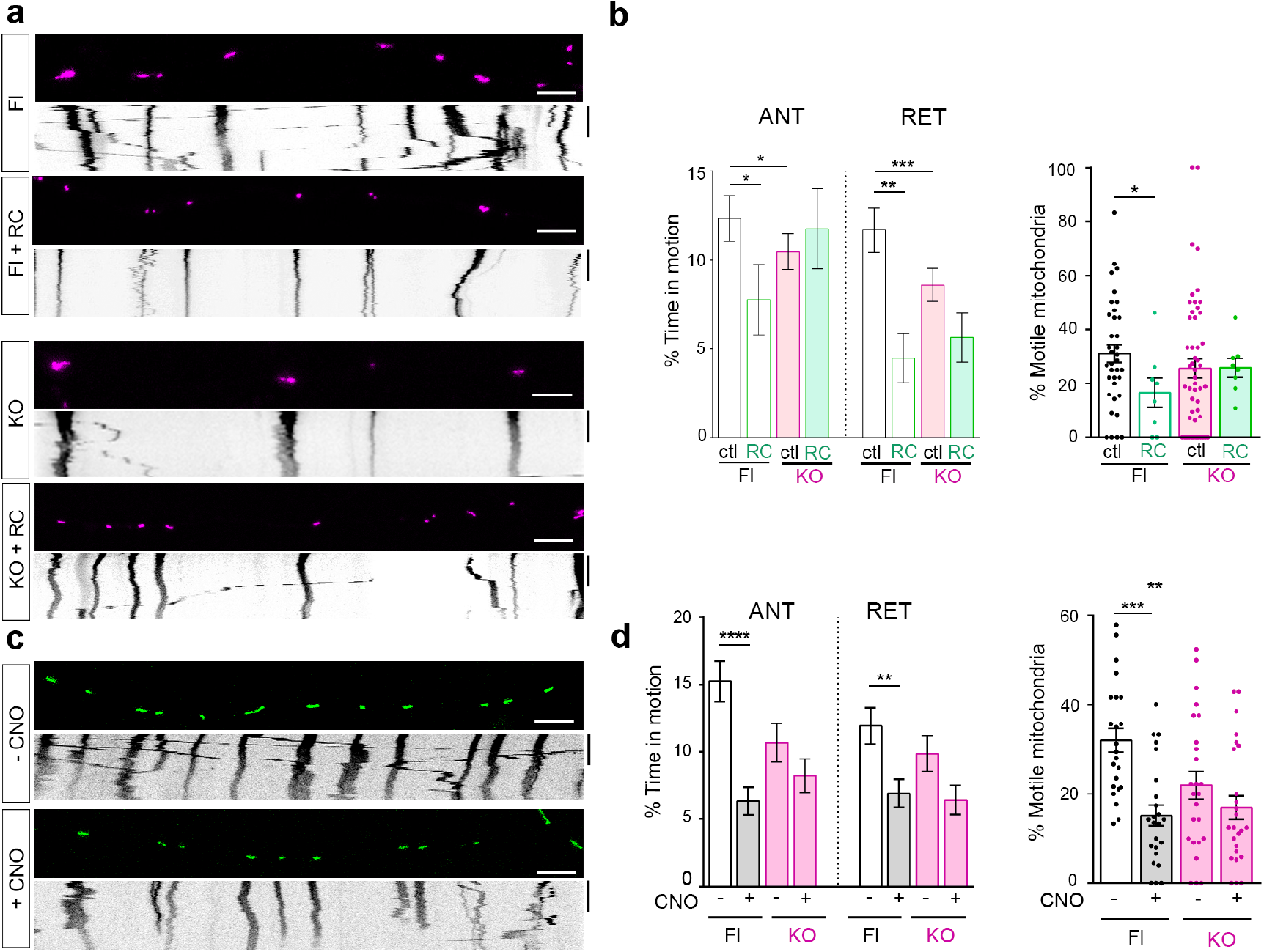
Fig. 4: The Gα_q_ effects on mitochondrial trafficking depend on Alex3. **a,** Images and kymographs from control (Fl) or *armcx3* KO (KO) mice axons expressing GFP and mitoDsRed in the presence or absence of Gα_q_R183C (RC). **b,**From kymographs as in (**a**), Gα_q_R183C decreased “% of time in motion” (TIM) and “% of motile mitochondria” (MM) in *armcx3* KO axons. n = 431 (Fl), 133 (FlRC), 685 (KO), and 154 (KORC) mitochondria from 36 (Fl), 8 (FlRC), 52 (KO), and 8 (KORC) independent axons. **c,** Images and kymographs from *armcx3* KO axons expressing mitoGFP and the Gα_q_-hM3D DREADD before (-CNO) or after (+CNO) the addition of 1 μM CNO. **d,**From kymographs as in (**c**), %TIM in the anterograde (ANT) and retrograde (RET) directions, and %MM. n = 426 (Fl-CNO), 413 (Fl+CNO), 346 (KO-CNO), and 359 (KO+CNO) mitochondria from n = 23 (Fl-CNO), 24 (Fl+CNO), 25 (KO-CNO), and 25 (KO+CNO) axons. Scale bar, 5 μm. Time bar, 300 sec. Data represent mean ± s.e.m. Statistical analyses: **b,d,** Kruskal-Wallis with Dunn correction **b**, %TIM ANT= 9.7 *P*= 0.0213, RET = 25.85 *P*< 0.0001; %MM = 25.40 *P*= 0.0001; **d**%TIM ANT = 2.541 *P*= 0.4679, RET = −3.031 *P*< 0.0001; %MM = 18.91 *P*= 0.0003. \**P*<0.05, \*\**P*<0.01, \*\*\**P*<0.001, \*\*\*\**P*< 0.0001.

We also analyzed the effects of activating the GPCR-specific Gα_q_ pathway on mitochondrial trafficking by expressing the DsRed-tagged hM3Dq DREADD-Gα_q_ receptor in *Nestin^Cre^/farmcx3* neurons. The effects against were not additive and the absence of Alex3 reduced the effects of CNO-induced DREADD activation on mitochondrial trafficking (Fig. 4c, d). Conversely, the increased mitochondrial trafficking observed upon Gα_q_ knock-down was mostly unaffected by the lack of Alex3 protein (Supplementary Fig. 4a, b), showing that this process is independent of Alex3.

Mitochondrial number and length were analyzed in axons of *Nestin^Cre^/farmcx3*neurons. Even though we found reduced numbers of mitochondria compared with control littermates, mitochondrial length was not altered (Supplementary Fig. 4c, d). Instead, expression of active Gα_q_ increased the number of mitochondria irrespective of Alex3 expression and decreased mitochondrial length only when Alex3 was present (Supplementary Fig. 4 e,f). Finally, a similar reduction in mitochondrial size was observed after ligand activation of DREADD-Gα_q_ (Supplementary Fig. 4g, h). Overall, these results suggest that the functions of Alex3 and Gα_q_ in controlling mitochondria motility and dynamics lie in the same pathway.

### Mitochondrial distribution depends on Alex3 and activation of Gα_q_

Defects in mitochondrial motility may cause altered distribution in dendrites and axons^37^. We thus performed a mitochondrial distribution analysis on hippocampal neurons (Fig. 5). The Mito80 value (ie, location of 80% of proximal mitochondria) revealed increased mitochondrial accumulation near the soma in *armcx3*-KO neurons, compared to control neurons (Fig. 5a-c). In contrast, expression of the active form of Gα_q_ produced a redistribution of mitochondria towards the periphery of dendrites (Fig. 5a, d). In line with this, active Gα_q_ expression prevented mitochondria from concentrating near the soma in the absence of Alex3 (Fig. 5a, e). Next, we measured the density of mitochondria in the cell bodies of cultured neurons. The mitochondrial accumulation seen in *armcx3*-KO neurons was partially reversed by expression of active Gα_q_ (Fig. 5f). In contrast, knock-down of Gα_q_ did not cause any significant changes in mitochondrial distribution in control or *αrmcx3*-KO neurons (Supplementary Fig. 5a, b).

**Fig. 5:**
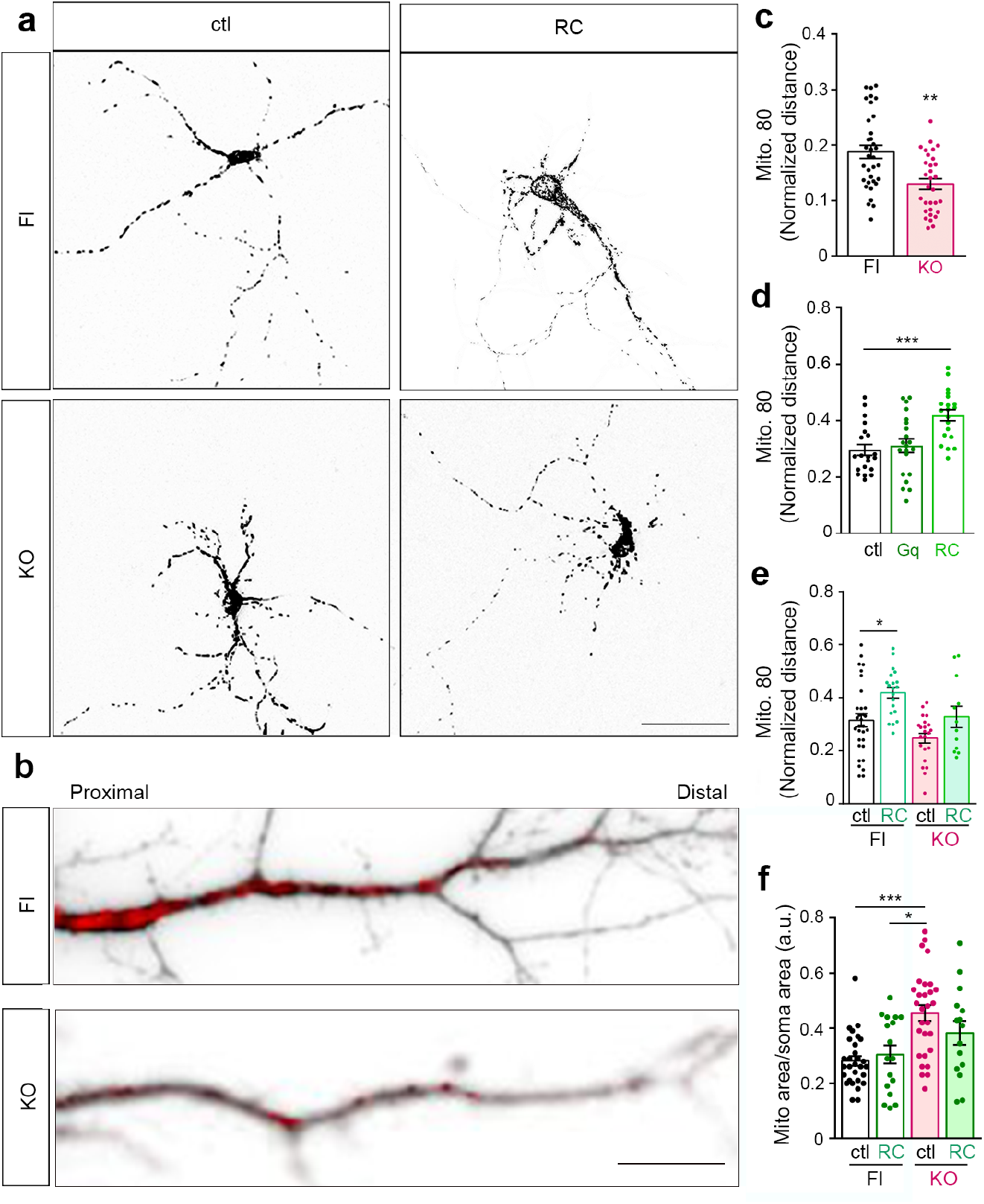
Mitochondrial distribution in dendrites depends on Alex3 and Gα_q_. **a**, Mitochondrial distribution in control (Fl) or *armcx3* KO (KO) neurons at 7 DIV expressing: GFP and mitoDsRed (ctl) in the presence or absence of Gα_q_R183C (RC). Scale bars, 50 μm. **b**, Dendritic processes (grey) showing mitochondrial distribution (red) in Fl and KO dendrites. Scale bars, 20 μm. **c, d, e**, Length-normalized Mito80 values (**c**) from Fl or KO neurons at 14 DIV. n = 34 (Fl) and 31 (KO) neurons; (**d**) from Fl neurons at 6-7 DIV expressing mitoDsRed and GFP without (ctl) or with Gα_q_ (Gq) or Gα_q_R183C (RC). n = 23 (ctl), 21 (Gq), and 13 (RC) neurons; (**e**) from Fl and *armcx3* KO neurons at 6-7 DIV expressing mitoDsRed without (ctl) or with Gα_q_R183C (RC). n = 54(Flctl), 20 (FlRC), 22 (KOctl), 12 (KORC). **f** The mitochondrial area in the soma of Fl or KO neurons expressing or not Gα_q_R183C was determined after subtraction of the nucleus area. n = 29 (Flctl), 17 (FlRC), 29 (KOctl) and 15 (KORC) somas. Data represent mean ± s.e.m from at least 3 independent experiments. Statistical analyses: **c,**unpaired two-tailed Student’s t-test with Welch correction T=3,8 df=61*P*= 0.0003; **d**, **e** two-way ANOVA with Bonferroni correction F(2,54) = 4.987 *P*= 0.0103; **e,** F(3,8) = 7.8 *P*= 0.0001**, f**, Kruskal-Wallis with Dunn correction for MA/SA= 20.00 *P*= 0.0002. *P*< 0.0001. * *P*< 0.05, ** *P*< 0.01, *** *P*< 0.001.

Altogether, these data indicate that absence of Alex3 results in an altered mitochondrial distribution in neurites concomitant to a mitochondrial accumulation in cell bodies. Activation of Gα_q_ partially reverts the effects of Alex3 depletion, indicating that Gα_q_ may function downstream of Alex3.

### Alex3 and Gα_q_ are required for proper dendritic arborisation

We next investigated the impact of Alex3-related mitochondrial alterations in dendritic morphogenesis and differentiation (Fig. 6). Compared with control neurons, we found an overall decrease in total dendritic length in *armcx3* KO neurons, along with a higher number of primary dendrites and a lower density of branching points, which correlated with an altered Sholl analysis (Fig. 6 a-c).

**Fig. 6:**
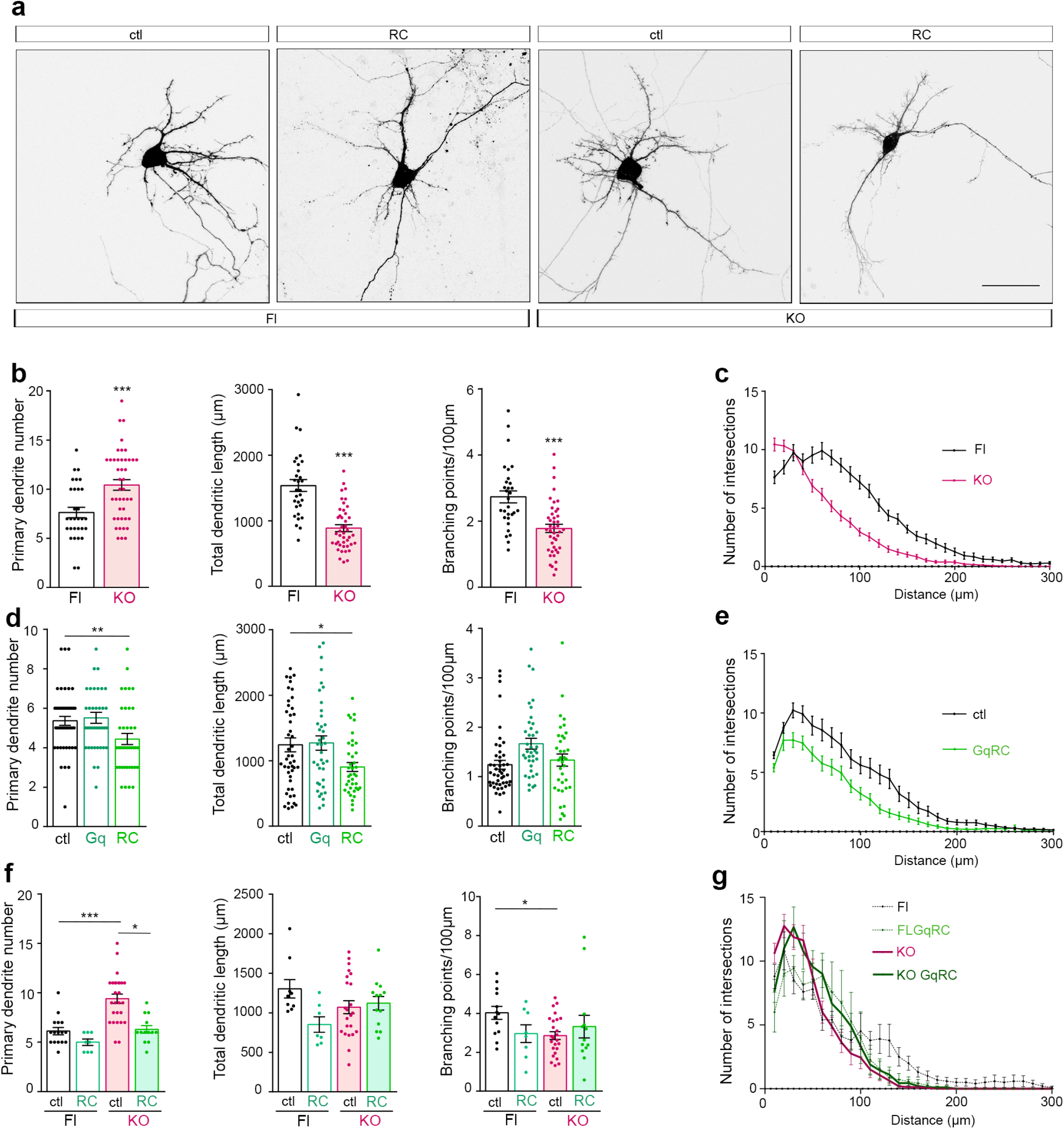
Alex3 and Gαq regulate dendritic arborization. **a,** Control (Fl) and *armcx3* KO (KO) neurons expressing GFP and mitoDsRed without (ctl) or with Gα_q_R183C (RC) imaged at 7 DIV. Scale bars, 50 μm. **b** and **c,** Fl and KO neurons expressing GFP at 14 DIV: average number of primary dendrites (PD, left), average length of dendritic tree (TD, middle), number of branch points normalized per total length of dendritic tree and expressed as a percentage (BP, right) and sholl analysis of branch points. n = 30 (Fl) and 42 (KO) neurons each for **b** and **c**. **d** and **e,** Neurons expressing mitoDsRed and GFP without (ctl) or with Gα_q_ (Gq) or Gα_q_R183C (RC) at 7 DIV, analyzed as in (**b**). Sholl analysis of branch points; n = 47 (ctl), 39 (Gq), and 39 (RC) neurons. **f** and **g,**Fl or KO neurons at 7 DIV expressing mitoDsRed and GFP without (ctl) or with GαqR183C (RC). n = 16 (Flctl), 8 (FlRC), 27 (KOctl), and 13 (KORC) neurons analyzed for primary dendrite number; n = 9 (Flctl), 7 (FlRC), 23 (KOctl), and 13 (KORC) neurons analyzed for total dendritic length; and n = 13 (Flctl), 8 (FlRC), 25 (KOctl), and 13 (KORC) neurons analyzed for the number of branch points. **g**, Sholl analysis of branch points. n = 32 (Flctl) and 39 (FlRC) neurons for (**d**); 9 (KOctl) and 43 (KORC) neurons for (**f**). Data represent mean ± s.e.m. Statistical analyses: **b,**unpaired two-tailed Student’s t-test with Welch correction, PD df = 69 *P*= 0.0004, TD df = 47 *P*= 0.0001, BP df = 54 *P*= 0.0001; **d,**Kruskal-Wallis with Dunn test statistic PD =10.57 *P*= 0.0051, TD =6.47 *P*= 0.0394, BP =11.29 *P*= 0.0035; **f,**Kruskal-Wallis with Dunn test statistic PD = 31.70 *P*< 0.0001, TD = 6.970 *P*= 0.0729, BD = 7.08 *P*= 0.0694. \**P*< 0.05, \*\**P*< 0.01, \*\*\**P*< 0.001.

Lower number and complexity of dendrites were also found in neurons expressing Gα_q_ R183C, whereas no reduction was found when Gα_q_ was expressed (Fig. 6a, d, e). Importantly, a gain of function of Gα_q_ (Gα_q_R183C) in *Nestin^Cre^/farmcx3* neurons largely reverted the alterations in primary dendrites observed upon Armcx3 deletion (Fig 6 f, g). Finally, we found that knock-down of Gα_q_ protein resulted in a decreased dendritic length and altered dendritic pattern, which was not observed when Gα_q_ was knocked down in *armcx3* KO neurons (Supplementary Fig. 6a-c). These results indicate that the dendritic effects observed by deletion of *armcx3* can be partially overcome by expressing the active form of Gα_q_ and, conversely, that Alex3 is required for most of the dendritic phenotypes driven by expression of active Gα_q_.

### Lack of Alex3 leads to decreased OXPHOS respiratory complex and to neuronal death

Western blotting experiments detected a significant decrease in the levels of mitochondrial respiratory complexes, supporting a reduction in OXPHOS activity in *Nestin^Cre^/farmcx3* mice (Supplementary Fig. 7a,b). These results were similar to those observed in *miro1/2* null mutants^38^. Next, we analyzed the contribution of ER stress response elements. The levels of pIRE1 and CHOP were reduced in *armcx3*-deficient brains, whereas eiF2α levels were unaltered (Supplementary Fig. 7c-k, m). In contrast, Bcl-2 and BIP protein levels did not show changes between control and *armcx3* KO mice (Supplementary Fig. 7l, n-p). These findings reflect alterations in the machinery that regulates mitochondrial respiration and ER stress response upon *armcx3* deletion.

Considering the findings above and the smaller brain size in *Nestin^Cre^/farmcx3* mice, we investigated whether *armcx3* deficiency affects neuronal cell death. To test this possibility, we quantified apoptotic (cleaved Caspase-3+) cells in several brain regions. Our results evidenced a marked increase in the number of apoptotic cells in *Nestin^Cre^/farmcx3* mice, which was especially conspicuous in the layers II-III of the cerebral cortex, the subiculum, the striatum, and the dentate gyrus (Fig. 7a-f). In addition, using the microglial marker Iba-1 we detected a dramatic and widespread increase in microgliosis across all regions studied in the absence of Alex3 protein (Fig. 7g-l). We conclude that deficiency of Alex3 in vivo results in increased neuronal death and neuroinflammation.

**Fig. 7:**
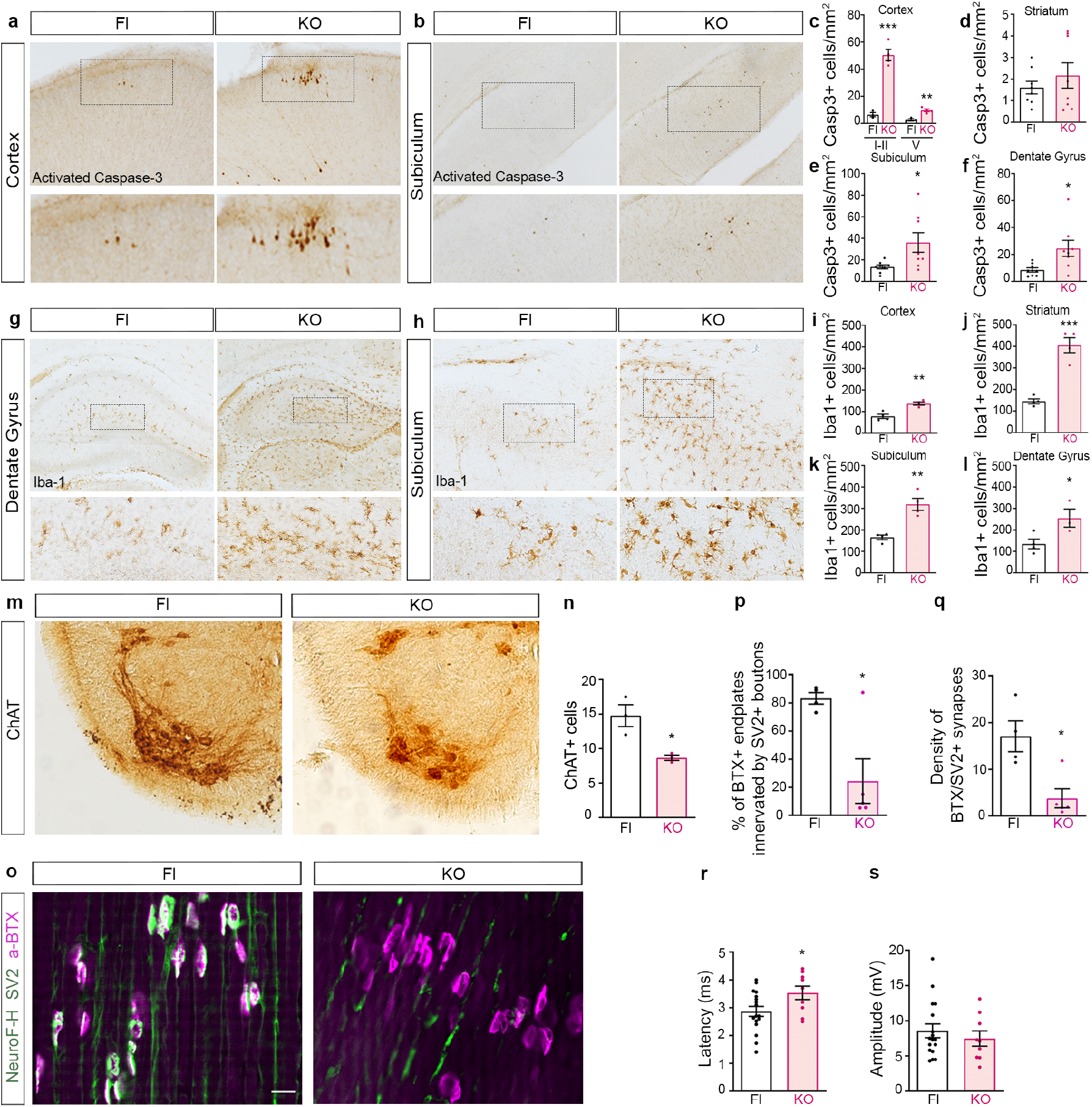
In vivo Alex3 deficiency increases cell death and microgliosis and leads to neuromuscular synaptic and electrophysiological deficits. **a-b**, Representative photomicrographs of activated caspase-3 in the cortex (**a**) and subiculum (**b**), evidencing an increased number of apoptotic cells in *armcx3* KO (KO) compared with control (Fl) mouse brains. **c-f,**Quantification of the density of activated caspase-3+ cells in layers I-II and V of the cortex (**c**), the striatum (**d**), the subiculum (**e**), and the dentate gyrus (**f**), further showing a general increase in the number of apoptotic cells in the absence of Alex3 in the brain. **g-h**, Representative photomicrographs of Iba-1 in the dentate gyrus (**g**) and subiculum (**h**) of Fl and KO brains. **i-l,**Quantification of microglial cell density in the cortex (**i**), the striatum (**j**), the subiculum (**k**), and the dentate gyrus (**l**), further showing a general increase in microgliosis in the absence of Alex3 in the brain. **m-n,** Choline acetyl-transferasepositive motor neurons in the thoracic ventral horn of the spinal cord; a decrease in number of motor neurons is noticeable in KO compared to Fl mice. **o,**Representative images of the neuromuscular junctions of the diaphragm muscle stained with rhodamine-conjugated α-bungarotoxin (BTX, magenta; post-synaptic acetylcholine receptors at endplates), and anti-neurofilament SMI-312 and anti-SV2 (green; axons and presynaptic terminals, respectively). **p-q,**Quantification of the % of BTX+ endplates innervated by SV2+ boutons (**p**) and the density of synapses double labeled with BTX and SV2 (**q**) in *armcx3* KO compared to Fl mice. **r-s,**In vivo motor nerve conduction analysis of peripheral nerve latency (stimulation at sciatic nerve and recording at gastrocnemius) (**r**) and amplitude (**s**), evidencing an impaired motor nerve conduction in KO compared to Fl mice. Data represent mean ±s.e.m. Statistical analyses: unpaired two-tailed Student’s t-test **c**, layer I-II, t = 8.515 df = 5 *P*= 0.0004, layer V, t = 5.136 df = 5 *P*= 0.0037; n = 3-4 mice per genotype; **d**, t = 0.8537 df = 14 *P*= 0.4076; n = 8 mice per genotype; **i**, t = 4.827 df = 6 *P*= 0.0029; **j**, t = 7.093 df = 6 *P*= 0.0004; **k**, t = 5.158 df = 6 *P*= 0.0021; **l**, t = 2.711 df = 5 *P*= 0.0422; n = 3-4 mice per genotype; **n**, t = 3.740 df = 4 *P*= 0.0201; n = 3 mice per genotype; **p**, t = 3.588 df = 7 *P*= 0.0089, n = 4-5 mice per genotype; **r**, t = 2.199 df = 24 *P*= 0.0377; **s**, t = 0.7039 df = 24 *P*= 0.4883; n = 9-17 mice per genotype;; unpaired two-tailed Student’s t-test with Welch’s correction **e**, t = 2.446 df = 7.563 *P*= 0.0419; **f**, t = 2.560 df = 8.019 *P*= 0.0336; n = 8 mice per genotype; **q**, t = 3.595 df = 4.528 *P*= 0.0185; n = 4-5 mice per genotype. \**P*< 0.05, \*\**P*< 0.01, \*\*\**P*<0.001.

### Deletion of *armcx3* results in decreased number of motor neurons and neuromuscular synapses and in physiological alterations

Because of the motor deficits observed in *Nestin^Cre^/farmcx3* mice, we then determined whether spinal cord motor neurons were affected in these animals by analyzing Chat (choline acetyltransferase) expression. We found a lower number of Chat+ motor neurons in the ventral horn of the spinal cord of *Nestin^Cre^/farmcx3* mice compared with littermate controls (Fig. 7m-n). To investigate whether neuromuscular synapses were altered, the neuromuscular junctions of diaphragm muscles from control and KO mice were visualized by labeling post-synaptic acetylcholine receptors at endplates with α-bungarotoxin, and axons and presynaptic terminals with Synaptobrevin 2 (SV2). Our results evidenced a dramatic decrease in the density of SV2-labeled endplates in *armcx3* KO mice compared with controls, hence indicating impairment in these neuromuscular synapses in the absence of Alex3 (Fig. 7o-q).

Finally, motor nerve conduction tests were performed at P5 to investigate physiological consequences of *armcx3* inactivation. After sciatic nerve stimulation, we found increased latencies of compound muscle action potentials (CMAPs) in *Nestin^Cre^/farmcx3* mice. In contrast, the amplitude of CMAPs did not differ between genotypes (Fig. 7r, s).

### Gα_q_ interacts with Alex3, Miro1 and Trak2

One of the characteristics of the G protein binding partners with armadillo domains is that they behave as chaperons for Gαsubunits ^39^. We found that levels of Gα_q_ were reduced in the brains of *Nestin^Cre^/farmcx3* mice compared to control littermates (Fig 8a). Interestingly, the levels of Alex3 were also markedly decreased in MEF cells knockout for Gα_q_/_11_^23^(Fig 8b).

**Fig. 8:**
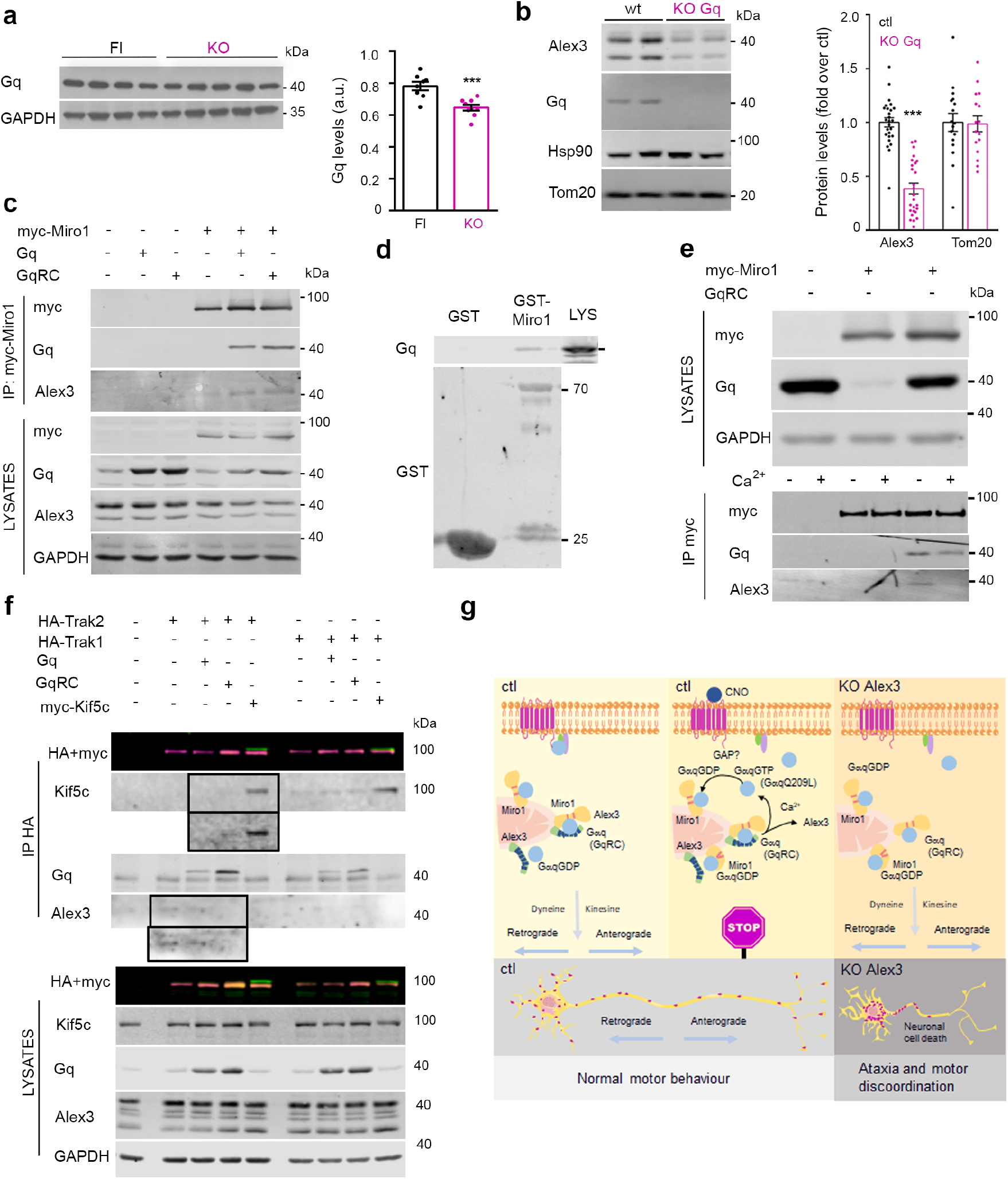
Gαq interacts with Alex3 and Miro1 and potentiates the binding to Trak2. **a**, Levels of Gα_q_ protein from P5 ccontrol (Fl) and *armcx3* KO (KO) mouse brains were analyzed by western blot (left). Standardization by GAPDH revealed a decrease in Gα_q_ expression in the absence of Alex3 (right). **b**, Levels of Alex3 and Tom20 proteins from WT and Gα_q_/_11_-/-(KO Gq) MEF lysates were analyzed by western blot (left). Standardization by Hsp90 unveiled a decrease in Alex3 expression in the absence of Gα_q_/_11_, while no significant variations in Tom20 levels were observed. **c**, Myc-Miro1 co-immunoprecipitated with Gα_q_ or Gα_q_R183C and endogenous Alex3 from extracts (700 μg) of HEK293. **d**, Gα_q_ binds Alex3 and Miro1 in the non-active state. Pull-down of GST-Miro1 from extracts of SHSY5Y cells followed by western blot revealed co-precipitation of endogenous Gα_q_. **e**, Myc-Miro1 immunoprecipitated with Gα_q_R183C in the presence of 2 mM calcium while Alex3 dissociated from the complex in HEK293 cells lysates. Lysates were split, and immunoprecipitation was carried out in the presence (+) or absence (-) of 2 mM calcium. **f**, Gα_q_R183C immunoprecipitates with HA-Trak1 or HA-Trak2 from lysates of HEK293 cells. Kif5c immunoprecipitates with Trak2 in the presence of Gα_q_R183C or with excess of Trak2 (high exposure). **g**, Diagram representing Gα_q_ signaling in the presence or absence of Alex3. Data represent mean ± s.e.m. of 3 independent experiments. Statistical analyses: **a,**unpaired two-tailed Student’s t-test with Welch correction *P*= 0.0012; **b**, one-way ANOVA with Bonferoni post hoc test F(3.01)= 30.5 \**P*< 0.05, \*\**P*< 0.01, \*\*\**P*< 0.001.

Given that Alex3/Gα_q_ protein complex regulates mitochondria motility, we investigated whether Gα_q_ is found in a complex with the mitochondria Rho-GTPase Miro1^14^. Immunoprecipitation of myc-Miro1 detected Gα_q_ and Gα_q_R183C together with endogenous Alex3 (Fig. 8c). Moreover, pull-down experiments of GST-Miro1 in SHSY5Y cell lysates showed that Gα_q_ interacts with both Miro1 and Alex3 (Fig. 8d). The presence of high (2 mM) calcium levels did not disrupt the interaction between Gα_q_ and Miro1, as did with the Alex3-Miro1 interaction (Fig 8e). Finally, we analyzed the interactions with Trak1, Trak2, and the motor protein Kif5c in the presence of active Gα_q_ and endogenous Alex3. Gα_q_ interacted with both Trak1 and Trak2. Endogenous Alex3 immunoprecipitated only with Trak2, as shown before^14^. Interestingly, the presence of active Gα_q_ protein potentiated the interaction between Trak2 and endogenous Kif5c, which is otherwise not present unless Kif5c is overexpressed (Fig. 8f, see overexposed images)^40^.

Together, these findings show that Alex3 and Gα_q_ proteins regulate each other’s levels, and more importantly, that Gα_q_ is part of the Miro/Trak2/Alex3 complex that regulates mitochondrial trafficking. Activation of the G protein would switch the complex towards a state of Trak2-driven mitochondrial movements that affect dendrite arborization and would contribute to calcium-induced mitochondrial arrest.

## Discussion

The present study provides evidence for a novel protein complex (Alex3/Gα_q_) that regulates mitochondrial trafficking and dynamics in neurons through its link to the well characterized Miro/Trak trafficking complex^4,6,14,40–44^. Mitochondrial Gα_q_ and Alex3 interact functionally in the regulation of important processes such as neuronal mitochondrial distribution and dendritic growth and complexity. Finally, because Gα_q_ signals via numerous important GPCR (eg, metabotropic glutamate receptors, muscarinic receptors, and many neuropeptide receptors), our findings suggest a novel molecular pathway by which a variety of extracellular signals may control mitochondrial dynamics and functions.

Our data reveal that stimulation of the GPCR-Gq pathway, or direct activation of Gα_q_, arrest mitochondrial motility. Since the PLCβ effector domain of Gα_q_ is not required for mitochondria arrest other pathways may be involved in that process. Our proteome analysis identified Alex3 as a partner of Gα_q_. We focused on this protein because it is a component of the mitochondria motility complex^14^ and it contains armadillo domains that interact to Gα subunits^39,45–47^ and GPCRs^12^. We provide evidence that Alex3 binds Gα_q_ through its armadillo domain but it does not act as an effector: Alex3 directly binds Gα_q_, predominantly its non-constitutively active form (Gα_q_Q209L), but can bind to Gα_q_-effector mutant^31,33^. In line with that, GRK2 expression, which blocks effector interactions, did not affect Alex3-Gα_q_ interaction^34^. We also showed that the phenotype of cell body mitochondrial aggregation observed upon *armcx3* deletion is diminished by expressing the active form of Gαq. Moreover, expression of active Gα_q_ partially reverts the effects of *armcx3* deletion on neuronal arborization, altogether suggesting that Gα_q_ effects lay downstream of Alex3 function.

Our results suggest that Alex3 regulates the complex established between Gα_q_ and Miro1 at mitochondria in a cyclic way: Alex3 would dissociate from Miro1 upon a rise in calcium levels^14^, in order to dissociate the whole complex from the microtubules, or it would change its interaction with the adaptor protein Trak2, in order to switch transport direction (Fig. 8g). Alex3 may have a role as scaffolding protein, or even as GEF or chaperone for Gα_q_^39,48^. In this cycle, Alex3 would potentiate the role of GPCRs to control mitochondrial movement^18,49^. However, depletion of Gα_q_ diminishes the levels of Alex3. Phosphorylation of Alex3 through the canonical PLCβ-PKC axis by Wnt leads to its degradation^35^, suggesting a regulatory feedback loop where Gα_q_ activation by the canonical pathway would control the levels of Alex3 protein. Overall, this additional regulation could contribute to the arrest of mitochondrial movement in response to calcium rises or to changes of directionality upon GPCR stimulation.

Other *GPRASP/ARMCX* family members may interact with G proteins, particularly when considering that Armc10 and Alex1 also associate with Miro1 and share some physiological functions with Alex3^17,35^, and that Armc10 was found in the Gα_q_ proteomic analysis (Fig. 2a). Moreover, GPRASP1 interacts with the delta opioid receptor and with the heterotrimeric Gα_s_ subunit^45^, and Armcx3 and Armc10 are related to the GPCR-effector AMPK^50,51^. We thus propose that members of the *GPRASP/ARMCX* family may be modulators of GPCR signaling, by interacting with either GPCRs *(GPRASP)* or Gα subunits (*ARMCX*), or by acting as their effectors.

No loss of function mutations and very low number of missense variants in *ARMCX3* have been identified in hemizygosis in human populations (Supplementary Table S1), suggesting that this gene may be intolerant to loss of function mutations. Besides, large duplications affecting *ARMCX3* are associated to schizophrenia and autistic disorders (ClinVar, nsv932069)^52^, whereas *ARMCX3* variants are associated to metabolic diseases^53^ and *ARMCX3* is the most prominent gene linked to sleep-disordered breathing hypoxia^54^(Supplementary Table 1). *GNAQ* (Gα_q_) mutations lead to motor and cognitive dysfunctions^24–26^, and cause seizures and mental retardation^55^, whereas mutations and variants in the functionally related genes *TRAK2* and *RHOT1/2*(Miro1/2) are associated to autism disorders and late-onset neurodegenerative diseases (Supplementary Table S1). Interestingly, Miro1 protein decreases mitophagy via Parkin^38,56^.

Several studies have stablished the relationship between altered dendritic development and defects in mitochondrial localization or trafficking machinery^37,57,58^. The present study highlights the relevance of Alex3 in distinct neural processes (dendritic formation, synaptogenesis and neuronal survival), which are likely involved in the occurrence of motor deficits. Although *miro1* ablation also drives loss of dendritic complexity, this happens without locomotion defects^37^, which is probably due to *miro1* ablation occurring postnatally.

Mitochondrial dysfunction and loss of dendritic and axonal complexity are hallmarks of many neurodegenerative disorders, including ataxic disorders and late-onset neurodegenerative diseases such as Alzheimer disease^7-–10,59–61^. Armc10 prevents Aβ-induced mitochondrial fission and neuronal death^16^. The present study shows that the Alex3/Gα_q_ complex is necessary for correct dendritic arborization and mitochondrial distribution in neurons. Moreover, the lack of Alex3 in vivo results in altered expression of mitochondrial respiratory complexes and ER stress response elements. *armcx3* KO mice display increased neuronal death in many CNS regions, including the spinal cord, concomitantly to decreased peripheral synaptic innervation and severe motor deficits, which is reminiscent of features observed in mitochondrial diseases^62^. Together, these findings lead us to propose a model in which disruption of the Alex3/Gα_q_ protein complex results in aberrant mitochondrial trafficking and distribution, altered respiratory function^23^, dysregulation of OXPHOS/ER stress response proteins, decreased neurite growth, and finally, neuronal cell death. It remains to be determined how extracellular signals regulate these processes through the GPCR/Gα_q_ pathway and, considering the above genetic data, the likely contribution of the Alex3/Gα_q_ complex in the pathogenesis of human neurodegenerative and psychiatric disorders linked to mitochondrial dysfunction.

## Data availability

All data are available in the main text or the Supplementary Information. All other data supporting the findings of this study are available from the corresponding authors on reasonable request. Inquiries concerning the Gq proteomics data should be directed to A.M.A., inquiries concerning *armcx3* KO model should be directed to E.S.

## Acknowledgements

We Sara E. Rubio for discussion and critical reading of the manuscript, and A. Reyes, B. Bouazza and A. Josa for help in some experiments. This work was funded by the Ministerio de Ciencia e Innovación (grants BFU2017-83379-R to AMA, SAF2016-76340R and PID2019-106764RB-C21 to ES, SAF2015-65633-R and RTI2018-099357-B-I00 to JAE, RTI2018-096386-B-I00 to XN, Severo Ochoa Excellence program to JAE and María de Maeztu Excellence program to ES) and Instituto de Salud Carlos III (CIBERNED to ES, CA, XN and ALdM; CIBERER to GM; CIBERFES to JAE, grant PI18/01066 to ALdM and a collaborative CIBERNED project to ES and ALdM); JAE is supported by the HFSP (RGP0016/2018) and the Pro CNIC Foundation. ALdM is supported by EiTB Maratoia, grant number BIO17/ND/023, and by Osasun Saila, Eusko Jaurlaritzako, grant number 2015111122. FJG-B was supported by Roche Stop Fuga de Cerebros (BIO19/ROCHE/017/BD). IIV was supported by a FI fellowship from AGAUR.

## Author contributions

IIV, SM and YM performed or participated in most experiments; AP and MH measured dendritic parameters; CB, JR and JAI conducted the proteome analysis and performed biochemical analyses of Alex3 and Gq; SM and YM generated the *armcx3* KO model; FU analyzed the spinal cord; CA, RDC-T and EV performed biochemical analyses in KO mice; ER provided support with live imaging acquisition and analysis; XN and JDV performed the in vivo electrophysiology experiments; FJG-B and ALdM performed the neuromuscular synapse analysis; GM performed genetic analysis; IIV, SM, YM, AMA and ES wrote the draft of the manuscript. All authors read and corrected the manuscript. YM, AMA and ES wrote the final manuscript. AMA and ES conceived and supervised the study.

