## Supplementary Methods for "A novel Alex3/Gα_q_ protein complex regulating mitochondrial dynamics, dendritic complexity, and neuronal survival"

### Materials and Methods

#### Plasmid vectors

pcDNA3.1(+) was purchased from Invitrogen. pEGFP was from BD Biosciences. Mitochondrial-targeted DsRed and GFP (pmitoDsRed and pmitoGFP) and pcDNA3KIF5c-myc were a gift from Antonio Zorzano (IRB Barcelona, Spain)[1] Alex3 expression vectors (pSecTag-myc-Alex3, pEGFP-Alex3, pE-Alex3-Nt-GFP, pE-Alex3-Ct-GFP, pE-Alex3-ΔCt-GFP, pE-Alex3-ΔNt-GFP, pE.Alex3-(1-106)-GFP and pE-Alex3 (1-45)-GFP) were described in Lopez-Domenech et al., 2012[2]. GST-Alex3 constructs were generated and subcloned into pGEX2T and pGEX4T vectors (Pharmacia). For that, the following oligonucleotides were used:

GST-Alex3-FL(30-379) and GST-Alex3 Nt(30-111), Fw: 5'-CGCGGATCCGGAAGAAAGCAGAACAAGGAG-3';

GST-Alex3-FL(30-379), GST-Alex3 Ct(111-379) and GST-Alex3-Ct2(273-379), Rv: 5'-GCGGAATTCCTTCCTGACTCTTTGGGAACATC-3';

GST-Alex3 Nt(30-111), Rv: 5'-GCGGAATTCCAGGAGAAGCCCTTTTCTGTAC-3';

GST-Alex3 Ct(111-379) and GST-Alex3 Ct1(111-279), Fw: 5'-CGCGGATCCCCTAATTCAGACGATACTGTTTTGTC-3';

GST-Alex3 Ct1(111-272), Rv: 5'-CCGCTCGAGCTTTCAGCCAGATTCAAAAGGAGC-3';

GST-Alex3-Ct2(273-379), Fw: 5'-CGCGGATCCAATCCAGCCATGACTAGAGAACTGC-3'.

pcDNA3-G$\alpha_q$, pcDNA3-G$\alpha_q$R183C and pcDNA3-GαqQ209L were cloned from pCIS vectors from Melvin I. Simon (Caltech)[3] and described in Johansson et al., 2005[4]. The constitutively active Gαq mutant protein that lacks the ability to interact with PLCβ (GαqQ209L/R256A/T257A) was provided by Richard Lin (Stony Brook University, New York, USA). pcDNAGαqR183C/R256A/T257A was provided by Federico Mayor (CBM, Spain). pCNA-myc-GRK2 was a gift from Jeffrey L. Benovic (The Kimmel Cancer Center, USA). pcDNA1-GFP-G$\alpha_q$R183C and pcDNA1-GqNt-GFP generously provided by Catherin Berlot (Yale University School of Medicine, USA) were described in Benincá et al., 2014[5]. pcDNA3.1-G$\beta_2$-FLAG and pcDNA3.1-G$\gamma_1$-HA were from the Missouri S&T cDNA Resource Center (www.cdna.org). pLKO1-shGq (sh1) sequence was obtained from MISSION Sigma (Functional Genomics Core Facility, IRB Barcelona) as described in

Benincá et al., [5]and subcloned into pLKO.3G vector to obtain pLKO.3G-shGq (sh2). pLKO.3G-shscr (scr) was from Merck-Sigma. pRK5Myc-Miro1, pGW1TRAK1-HA, pGW1TRAK2-HA, pEGFP-synaptophysin and pCAGIRES-mtDsRed were kindly provided by Josef Kittler (University College of London, United Kingdom). For the generation of pIRES-mtDsRed-Gq, pcDNA3-G$\alpha_q$ was subcloned into pCAGIRES-mtDsRed vector using the primers 5’-CCGGGCTAGCATGACTCTGGAGTCCATCATGGCG-3’ and 5’-GGCCCGATCGTTAGACCAGATTGTACTCCTTCAGGTTC-3’. pIRES-mtDsRed-GqR183C was generated by single-point mutation from pIRES-mtDsRed-Gq using the QuickChange II site-directed mutagenesis Kit (Agilent) according to the manufacturer’s guidelines and using the primers 5’-GACGTGCTTAGAGTTTGTGTCCCCACTACAGGGA-3’ and 5’-TCCCTGTAGTGGGGACACAAACTCTAAGCACGTC-3’. pcDNA3-G$\alpha_q$R183C/R256A/T257A and pcDNA1-GFP-G$\alpha_q$R183C/R256A/T257A were generated by single-point mutation in two different reactions from pcDNA3-G$\alpha_q$R183C and pcDNA3-GFP-G$\alpha_q$R183C, respectively, using the QuickChange II site-directed mutagenesis Kit (Agilent) with the following primers: for PCR1,
Fw: 5’-GAGCAAAGCACTCTTTGCAACAATTATCACCTACC-3’;
Rv: 5’-GGGGTAGGTGATAATTGTTGCAAAGACTGCTTTGCT-3’; and for PCR2,
Fw: 5’-GCAAAGCACTCTTTGCAGCAATTATCACCTACCCC-3’;
Rv: 5’-GGGGTCGGTGATAATTGCTGCAAAGAGTGCTTTGC-3’.
pAAV-hSyn-hM3D(Gq)-mCherry was a gift from Bryan Roth (Addgene plasmid # 50474 ; http://n2t.net/addgene:50474 ; RRID:Addgene_50474) .

#### Antibodies

Primary antibodies and dilutions used were: Alex3 (25705-1-AP, ProteinTech, 1:1000), β-tubulin (MMS-431P, Covance, 1:10000 and T2200 Merck-Sigma, 1:10000), GFP (A11122, Invitrogen, 1:1000 and gta-20, Chromotek, 1:200), DsRed (632496, Clontech, 1:2000), GST (SAB4200237, Merck-Sigma, 1:2000), myc (05-724, Millipore, 1:2000; M4439, Merck-Sigma, WB:1:2000, IF: 1:200; and yta-20, Chromotek, 1:200), HA (11867423001, Roche, 1:2000 and A2095, Merck-Sigma, 1:25), FLAG (F1804-50UG, Merck-Sigma, 1:1000), Gαq (G7, sc-136181, Santa Cruz, 1:1000; E17, sc-393, Santa Cruz, 1:1000; C19, sc-392, Santa Cruz, 1:1000; and 612704, BD, 1:1000), Miro1 (HPA010687, Merck-Sigma, 1:1000), GAPDH (14C10, Cell Signaling Technology,

1:1000), HSP90 (610418, BD, 1:2000), TOM20 ((sc-17764, Santa Cruz, 1:1000 and sc-11415, Santa Cruz, 1:1000)), pIRE(pSer724) (NB-100-2323, Novus, 1:0000), Bcl2 (50E3) (2870, Cell Signaling Technology, 1:1000), BIP (3183, Cell Signaling Technology, 1:1000), CHOP (L63F7) (2895, Cell Signaling Technology, 1:1000), eI2a (9722, Cell Signaling Technology, 1:1000), peI2a(ser51) (9721, Cell Signaling Technology, 1:1000), IRE1α(14C10) (3294, Cell Signaling Technology, 1:1000), OXPHOS ( 110413, Abcam, 1:1000), Iba-1 (019-19741, Wako, 1:1000), cleaved Caspase-3 (9664S, Cell Signaling Technology, 1:1000), ChAT (AB144P, Merck-Sigma, 1:100), Cux1 (SC-13024, Santa Cruz Biotechnology, 1:250), (CTIP2 (ab18465, Abcam, 1:500), calbindin, (300, Swant, 1:5000), SV2A (SV2, DSHB, 1:100), neurofilament H (clone SMI 312, 801701, BioLegend, 1:50), tetramethylrhodamine-conjugated α-Bungarotoxin (T0195, Merck-Sigma, 2 µg/ml).

Secondary antibodies and dilutions used were: HRP anti-mouse, polyclonal goat (Dako, 1:2000), HRP anti-rabbit, polyclonal swine (Dako 1:2000), IRDye 680 anti-mouse (925-68070, LI-COR, 1:20.000); IRDye 800 anti-mouse (925-32210, LI-COR, 1:20.000); IRDye 680 anti-rabbit (926-68073, LI-COR, 1:20.000); IRDye 800 anti-rabbit (925-32211, LI-COR, 1:20.000) and IRDye 680 anti-rat (925-32211, LI-COR, 1:20.000). For immunofluorescence in tissue or cell cultures, Alexa Fluor 488 anti-mouse (A21202), Alexa Fluor 568 anti-mouse (A11004), Alexa Fluor 647 anti-mouse (A21236), Alexa Fluor 488 anti-rabbit (A21206) and Alexa Fluor 647 anti-rabbit (A21244), all from Thermo Scientific, were used at 1:500 (tissue) or 1:2000 (cell cultures) dilutions.

#### Animals

Mice were bred, housed and studied in the animal research facilities at the University of Barcelona. Animals were provided with food and water *ad libitum* and maintained in a temperature-controlled environment in a 12/12 h light-dark cycle. All the experiments involving animals were performed in accordance with the European Directive 2010/63/EU and the National Institute of Health guidelines for the care and use of laboratory animals. All the experiments involving animals were approved by the local ethics committee for animal experimentation of the University of Barcelona (CEEA, Barcelona, Spain) with the exception of *in vivo* electrophysiology procedures, which were approved by the ethics committee of the Autonomous University of Barcelona (Barcelona, Spain).

#### Cell culture and transfections

HEK293 (ATCC), COS-1 (ATCC), Murine embryonic fibroblast (MEF), and MEF cells knock-out for $G\alpha_q$ and $G\alpha_{11}$ (a gift form Stephan Offermmans, Max-Planck-Institute for Heart and Lung Research, Germany) described before[5] were cultured in DMEM medium (GIBCO) supplemented with 10% (v/v) fetal bovine serum and 2 mM glutamine, and maintained at 37ºC in a 5% $CO_2$ incubator. SHSY5Y (ATCC) cells were cultured in DMEM/F12 medium (GIBCO). Cells were plated 24 h before transfection with FUGENE6 (Promega) or Lipofectamine 2000 (Life Technologies) according to the manufacturer's guidelines.

Mouse hippocampal cultures were obtained from E15 mouse embryos (OF1, Iffra Credo, Lyon, France) as described earlier[2]. Hippocampal cultures at 4 DIV in Neurobasal medium containing 0.59% D-glucose were transfected with Lipofectamine 2000 (Life Technologies) according to the manufacturer's guidelines. Essentially, 2 µg of DNA were complexed with 2 µl of Lipofectamine 2000 for 20 min before adding the mix to the media, then medium was replaced after 30 min, and cells further cultured 1-3 more days. The combinations of plasmids used for neuronal transfections were: a/ pmitoGFP and pAAV-hSyn-hM3D(Gq)-mCherry (for stimulation of DREADDS); b/ pEGFP and pIRES-containing vectors expressing mitoDsRed (for $G\alpha_q$ expression experiments with bicistronic vectors); c/ pmitoDsRed and pEGFP- or pEGFP-Gq-containing vectors; or d/ pmitoDsRed and pLKO.3G-sh scr (scr), or pmitoDsRed, pEGFP and pLKO1-shGq (sh1 Gq), or pmitoDsRed and pLKO.3G shGq (sh2 Gq).

#### Immunoprecipitation and western blot

For co-immunoprecipitation experiments, whole cell lysates in lysis buffer (50 mM Tris-HCl pH7.5, 150 mM NaCl, 5 mM EDTA, 1% Triton X-100, 10% Glycerol, 10 mM NaF, 50 mM $Na_2H_2P_2O_7$, 1.5 mM $MgCl_2$ with protease inhibitors) were centrifuged and supernatants were incubated under gentle rocking with a specific antibody (1-5 µg) and 1.5 µg of IgG-free bovine serum albumin (BSA) (Sigma) for 3 h or overnight at 4°C.

Antibodies were bound to beads by addition of 10-20 µg protein G or protein A Sepharose beads (Merck-Sigma) to each sample. When using a bead-TRAP-conjugated antibody (anti-myc and anti-GFP TRAP, Chromotek) or antibodies conjugated to protein G agarose beads (anti-HA sepharose beads, Merck-Sigma), 2-10 µl of antibody-beads were added per sample and incubated for 1-2 h at 4°C. The BioRad protein assay (BioRad) was used for protein quantification on lysates. Samples were rinsed with wash buffer (10 mM Tris HCl pH 8.0, 300 mM NaCl, 1 mM EDTA, 1 mM EGTA, 1% Triton X-100, 0.5% NP-40, 1.5 µM $MgCl_2$) and resolved by SDS-PAGE previous to transferring them to Immobilon-FL membranes (Millipore). Membranes were incubated with anti-mouse or anti-rabbit antibodies (IRDye 680 and IRDye 800) diluted 1:20.000 in Odyssey Blocking buffer (LI-COR). Labeling was visualized with an Odyssey infrared scanner (LI-COR). For quantification, Image Studio Software (LI-COR) was used to determine the integrated optic density (IOD); values were normalized to the loading control.

#### GST pull-down assay

GST and GST-fusion proteins were transformed into *E.Coli* strain BL21 and purified with glutathione beads (GE Healthcare). GST-proteins in 50 mM Tris-HCl pH 7.5, 150 mM NaCl, 1 mM NP-40, 0.25% deoxycholate, 1 mM EGTA pH 8.0, 1 mM NaF were incubated with lysates of SHSY5Y cells (around 1.5 mg of protein) or HEK293T cells transiently transfected with the plasmids indicated in each figure (700 µg of protein lysates) under gentle rotation for 1 h at 4°C. Samples were rinsed 4 times with wash buffer, analyzed by SDS-PAGE, and visualized by either ODYSSEY analysis or ponceau (Merck-Sigma) staining. In pull-down experiments with purified components, 10 µg of the bead-conjugated GST-Alex3 protein were incubated with 10 ng of purified Gαq in 1 ml of incubation buffer (100 mM NaCl, 20 mM HEPES, 2 mM $MgCl_2$, 0.5 mM EDTA,1 mM DTT).

#### Imaging of fixed cells

Cells were seeded on 35-mm, glass-bottom dishes (ibidi) or on 1.5-mm coverslips, fixed with 4% Paraformaldehyde (PFA) in PBS for 15 min, washed three times with PBS, and blocked in a solution containing 1% BSA and 0.1% Triton in PBS. Cells were incubated for 1 h with primary antibodies at the corresponding dilution in blocking buffer, then washed 3-5 times with PBS before being incubated with the corresponding secondary antibody at room temperature, washed 4 times in PBS, and mounted on Prolong Diamond (Invitrogen). All secondary antibodies (Alexa Fluor 488-, Alexa Fluor 555-, and Alexa Fluor 657-conjugated, Life Technologies) were used at a 1:2000 dilution. Otherwise, neurons were incubated with 1 µM MitotrakerGreen FM (Thermo Scientific) for 45 min at 37°C before fixation as explained above. Images of fixed cultures were taken on a LSM700 confocal microscope (Zeiss) using a 40x (NA 1.3) or 63x oil objective (NA 1.4).

#### In vivo imaging of hippocampal neurons

In vivo imaging experiments were performed at 37°C in an atmosphere of 5% $CO_2$ with a LSM780 confocal microscope (Zeiss) equipped with 40x (NA 1.3) and 63x (NA 1.4) oil objectives. All electronics were controlled through the ZEN software (Zeiss). mitoDsRed/mitoGFP-labelled mitochondria in axons were live imaged 1-2 days after transfection (4-5 DIV cultures). In mitochondrial tracking experiments, a Z stack of 7 images of the neuron was acquired from at a 2048x2048 pixel resolution for subsequent analysis of morphology and mitochondrial distribution. An axonal segment located approximately 90 to 160 µm distal to the soma was selected for live imaging. Z stacks of 7 images from the axonal region were taken every 6 s during 10 or 15 min using the mitoDsRed channel with a 800x100 pixel resolution and an extra 2x digital zoom. For studies where DREADD receptors or GFP-synaptophysin were expressed, mitochondria or synaptophysin-containing vesicles were imaged using the GFP channel. Movies were processed using ImageJ software (http://imagej.nih.gov/ij/), and kymographs were generated by tracing axons in their z-projections. In kymographs, straight vertical lines were considered as static mitochondria, and motile mitochondria (non-straight vertical lines) were traced to evaluate their motility and directionality[6–8]. The percentage of time in motion was calculated as the percentage of time a given mitochondrion (static or motile) spent moving at speed over 0,0083 µm/s towards the anterograde or retrograde direction and represented as an average. The percentage of motile mitochondria represents the relation between number of motile and static mitochondria for each condition. In control neurons, a similar fraction of anterograde and retrograde movement is normally observed. The percentage of mitochondria in movement (20-30%) and the average velocity (between 0.3 and 0.7 µm/sec) also agreed with previous studies, which supports the validity of the methodology used in this study[9].

#### Neuronal and cell treatments

In live-imaging experiments using the G$\alpha_q$ specific inhibitor, neurons transfected with the plasmids of interest were incubated with 10 µM YM-254890 (Focus Biomolecules) during 30 min prior to imaging, as previously reported[10,11]. In GPCR-activation assays, neurons were co-transfected with mitoGFP along with a Cherry-tagged hM3D (Gq) DREADD receptor[12]. 24h after transfection, axons of DREADD-expressing neurons were imaged before and 15 min after the addition of 1 µM clozapine-N-oxide (Tocris). For determination of fluctuations on mitochondrial membrane potential, HEK293 cells expressing IRES vectors (with mitoDsRed) were incubated with 100 nM Mitotraker green (Thermo Scientific) for 30 min 24 h after transfection, followed by incubation with 10 µM valinomycin for 15 min before *in vivo* imaging. Live cells expressing mitoDsRed (IRES-mitoDsRed vectors with and without Gq) and MitotrakerGreen were imaged at 37°C in an atmosphere of 5% $CO_2$ with a LSM780 confocal microscope (Zeiss) equipped with a 63x (NA 1.4) oil objective. Confocal images in the green and red channels were acquired, and the intensity of green fluorescence was analyzed with the NeuronJ plugin from the ImageJ software. The fluorescence intensity density was determined for each cell and averaged.

#### Generation of a CNS-specific *Nestin$^{Cre}$/farmcx3* KO mice

To generate a CNS-specific Alex3 conditional knockout line, floxed*armcx3 (farmcx3)*+/+ female mice[13] were mated with *Nestin-Cre*+/- male mice[14], which express the Cre recombinase under the control of the Nestin promoter (active in neural and glial precursor cells). The following experimental groups were obtained: *farmcx3+/-;NestinCre-/-* (Fl control females, 25%), *farmcx3+/Y;NestinCre-/-* (Fl control males,

25%), *farmcx3+/-;NestinCre+/-* (heterozygous [Het] females, 25%) and *farmcx3+/Y;NestinCre+/-* (KO males, 25%). Unless otherwise stated, *farmcx3+/Y;NestinCre-/-* males and *farmcx3+/-;NestinCre-/-* females were used indistinctively as controls. The mating day was considered as embryonic day 0 (E0) and the day of birth as postnatal day 0 (P0). Mice at the following developmental stages were studied: E14, P5-P10.

#### Genotyping

Control, heterozygous and KO mice were genotyped using two complementary PCRs on genomic DNA extracted from tails. For the detection of floxedAlex3, the following primers were used: *Armcx3 S1F*, 5´-GGGGCGGTGGGCAGGATGACAG-3´; *Armcx3 S4*F, 5´-AAGTTCTAGGAATCGAGAGCC-3´; and *Armcx3 S*, 5´-ATCATTTCCCCTTGACTCTGG-3´. For the detection of the NestinCre allele, the following primers were used: WT-Cre Fw, 5´-CTAGGCCACGAATTGAAAGATCT-3´; WT-Cre Rv, 5´-GTAGGTGGAAATTCTAGCATCATCC-3´; Cre Fw, 5´-GCGGTCTGGCAGTAAAAACTATC-3´; and Cre Rv, 5´-GTGAAACAGCATTGCTGTCACTT-3´.

#### Behavioral phenotyping

Behavioral tests were performed in mice to assess their general neuromuscular function, body muscle strength, and posture. Mice were analyzed at P6-8. Phenotyping included body weight measurement, hind limb clasping tube test, hind limb suspension test, and kyphosis evaluation. Performance for each measure was recorded on a scale of 0-4 as described in El-Khodor et al., 2008 and Guyenet et al., 2010[15,16].

#### Tissue processing and histology

Postnatal mice were anesthetized with isoflurane and perfused with 4% PFA in 0.1 M Phosphate Buffer (PB). Brains and spinal cords were carefully extracted, post-fixed overnight with 4% PFA in PB, cryoprotected with 30% sucrose in PBS, and frozen at -42°C in isopentane. Frozen brain and cerebellum samples were sectioned in 50-µm coronal or sagittal sections, respectively, using a freezing microtome (Leica). Spinal cord samples were sectioned in a cryostat (Leica), either at 16 µm and collected in adhesive slides, or at 30 µm and stored free floating. All free-floating sections were collected in cryoprotectant solution (85% glycerol, 100% ethylene glycol, 0.1M PBS) and kept at -20°C until use. In addition, intact (whole-mount) diaphragms were also obtained to assess neuromuscular synapses.

The overall histological structure, width, and length of several brain regions were assessed using Nissl and/or DAPI staining.

For chromogenic immunodetection of antigens, sections were incubated with 10% methanol and 3% $H_2O_2$ in PB, to inactivate endogenous peroxidase activity, and then blocked for 2 h with 10% of either normal goat serum (NGS) or normal horse serum (NHS), 0.3 % Triton-X-100, and anti-mouse IgG F(ab')2 fragment when needed (Jackson ImmunoResearch, 1:300), in 0.2 % gelatin-PBS. Primary antibodies were incubated overnight at 4 °C in 5% NGS/NHS, 0.3% Triton-X-100 in PBS-0.2% gelatin. Sequential incubation with biotinylated secondary antibodies (1:200; 2 h at room temperature (RT)) and streptavidin-HRP (1:400; 2 h at RT), each of them diluted in 5% NGS/NHS and 0.3% Triton-X-100 in PBS-0.2% gelatin, was performed. Bound antibodies were visualized using 0.03% diaminobenzidine (DAB) and 0.006% $H_2O_2$ in PB as peroxidase substrates. Sections were mounted on gelatinized slides, dehydrated, and finally mounted in Eukitt medium (Merck-Sigma).

For immunofluorescence, frozen brain sections were permeabilized and incubated for 2 h at RT with 0.2% Triton-X-100, 1% BSA, 0.2M Glycine, and F(ab')2 fragment anti-mouse IgG (1:300) when needed, in 0.2% gelatin-PBS. For CTIP2 detection, heat-mediated antigen retrieval was performed before the blocking. The same solution was used for primary antibody incubation, performed overnight at 4°C, and secondary antibody incubation (Alexa Fluor, Invitrogen, 1:500), performed for 2h at RT. Nuclei were stained using DAPI (1:200), and sections were mounted in Mowiol medium (Calbiochem).

For neuromuscular synapses staining, samples were permeabilized in 0.5% Triton-X-100 in PBS and blocked in 10% donkey serum in 0.1M PBS for 1 h at RT before incubation with primary antibodies directed against synaptic vesicle protein SV2A and neurofilament H (clone SMI 312) for 2 days at 4°C. After several washes in 0.5% Triton-X-100, 1% donkey serum in 0.1M PBS, samples were incubated overnight with Alexa Fluor 488 donkey anti-mouse secondary antibody and tetramethylrhodamine-conjugated α-Bungarotoxin in 0.1M PBS. Samples were washed in 0.5% Triton-X-100, 1% donkey serum in 0.1M PBS and whole-mounted in Prolong Gold (Invitrogen) or Mowiol medium (Merck-Sigma).

#### Image acquisition and analysis of *Nestin$^{Cre}$/farmcx3* KO tissue

Bright field images were captured with a digital DP72 camera (Olympus) attached to an ECLIPSE 600 (Nikon) optical microscope with Cell T^ software and a NanoZoomer 2.0-HT (Hamamatsu Photonics). Immunofluorescence images were acquired with either an ECLIPSE E1000 (Nikon) optical microscope combined with a Cool SNAP camera, or with SPE (Leica) or TCS SP2 (Leica) confocal microscopes.

Low magnification (2x) images of DAPI-stained sections were used to measure the maximum width and length of the brain hemispheres at bregma positions ranging from +4.11 mm to +4.95 mm (according to Franklin and Paxinos, 1997) using Image J (FIJI) software. Additionally, the width of the primary somatosensory cortex was also measured using 10x images of DAPI-stained sections. The number of P5 brains analyzed ranged between 3 and 5 per genotype.

Images of Nissl-stained cerebellum sections from control and KO P10 mice were used to measure the maximum length of lobule IIc and IIIa (both in the same histological section), and lobule III and IV (both in the same histological section). One section per animal and 4-6 animals per genotype were analyzed for each of the parameters measured.

To determine the number and morphology of Purkinje cells, cerebellum sections from P10 mice were stained with anti-calbindin antibody. The number, body size and dendritic tree width (as a measure of the molecular layer width) of Purkinje cells were analyzed. Between 1 and 3 sections per animal and 5 animals per genotype were analyzed for each of the parameters measured.

To assess cell death and microgliosis, P5 brain sections were stained with antibodies against either the cleaved form of Caspase-3 or Iba-1, respectively. Several brain regions were assessed, including cortex (layers II-III and layer V), striatum, subiculum, and dentate gyrus. In each region, the number of cells with positive staining within a defined area was counted and expressed as number of positive cells per $mm^2$. The number of mice analyzed ranged between 3 and 8 per genotype.

The number of spinal ventral horn motor neurons was determined by counting the choline acetyltransferase (Chat)-positive cells that contained a clearly visible nucleus in serial P5 thoracic spinal cord sections. At least 3 sections per animal and 3 animals per genotype were analyzed.

Intact (whole-mount) diaphragms from 4-5 P6 mice per genotype were used to image neuromuscular synapses. Neuromuscular synapses were labeled using established markers of presynaptic nerve terminals (neurofilament H and synaptic vesicle protein SV2A) and postsynaptic motor endplates (α-bungarotoxin, which binds to acetylcholine receptors on the muscle fibers). Muscle preparations were visualized using a laser scanning confocal microscope LSM900 and Zen software (Zeiss). Neuromuscular synapses from 3 regions across the diaphragm were assessed in each preparation. For presynaptic analyses, endplates were categorized as either vacant (no neurofilament H/SV2 overlapping the endplate) or fully occupied (neurofilament H/SV2 overlapping more than 80% of the endplate). The synaptic density and the percentage of fully occupied endplates were calculated.

#### Motor nerve conduction tests

Motor nerve conduction tests were performed stimulating the sciatic nerve through a pair of needle electrodes placed at the sciatic notch. The compound muscle action potential (CMAP) was recorded from the gastrocnemius muscle (GM) with microneedle electrodes[17]. Potentials were amplified and displayed on a digital oscilloscope (Tektronix 450S; Tektronix). During the tests, the mouse body temperature was maintained between 34°C and 36°C on a heating pad.

#### Sholl and mitochondrial Sholl analysis

For arborization analyses, samples and procedures were as follows: fixed neuronal cultures permeabilized with 0.1% Triton X-100 in PBS for 5 min, blocked with 10% serum in PBS, and incubated in 5% serum overnight with anti-GFP antibody, of which images were taken with a LSM780 (Zeiss) confocal microscope equipped with a 20x objective. Images from live neuronal cultures were acquired on a LSM780 (Zeiss) upright confocal microscope equipped with a 63x (NA 1.4) or 40X (NA 1.3) objective (2048x2048 pixel resolution).

Some images were stitched together using Zen software (Zeiss) to allow visualization of the whole dendritic arbor. Dendrites were traced using NeuronJ pluguin from ImageJ software for quantification of dendrite number and length, number of branch points and Sholl analysis. Mitochondrial Sholl analysis was performed using a custom ImageJ plugin as previously described[7], which quantified the amount of mitoDsRed2 pixels within shells radiating out from the soma at one-pixel intervals. The density of the mitochondria in the soma (mito area/soma area) was calculated utilizing the ImageJ intensity density of mitoDsRed pixels from the soma of imaged neurons after subtraction of the intensity density of the nuclei in the red channel and averaged.

#### Mitochondrial number and length

Mitochondrial number and length were determined from axonal proximal segments of 25-40 neurons per condition. Live neuronal cultures expressing mitoDsRed were imaged with a LSM780 (Zeiss) confocal microscope equipped with a 40x (NA 1.3) and 63x (NA 1.4) oil objectives. Confocal images of the red (mtDsred) channel were acquired, then analyzed with the NeuronJ plugin of ImageJ software. Essentially, number of mitochondria within the axon was quantified manually and standardized to the length of the axonal section imaged, with axonal and mitochondrial length determined using the segmented line tool.

#### Tissue processing and western blot experiments in *Nestin*$^{Cre}$*/farmcx3* mice

Brains and spinal cords were rapidly dissected and isolated from E14 and P5-7 mice, frozen in liquid nitrogen, and stored at -80 °C before processing. Frozen tissue was lysed using a power homogenizer (Polytron) in 10 volumes of RIPA lysis buffer (50 mM Tris-HCl, 150 mM NaCl, 1 mM EDTA, 0.5% Sodium deoxycholate, 0.1% SDS, 1% Triton X-100) containing CompleteMini protease inhibitor cocktail (Roche) and phosphatase inhibitors (10 mM tetra-sodium pyrophosphate, 200µM sodium orthovanadate, and 10 mM sodium fluoride). After centrifugation of the lysates, supernatants were collected and stored at -80 °C. BCA protein assay (Thermo Scientific) was used for protein quantification.

Brain and spinal cord protein samples were resolved in SDS-polyacrylamide gels and transferred onto nitrocellulose membranes. Membranes were then blocked in 5% non-fat milk powder in TBST (10 mM Tris pH 7.4, 140 mM NaCl, 1% Tween-20) and incubated with the primary antibodies overnight at 4°C. For chemiluminescent detection, membranes were incubated with secondary HRP-labeled antibodies (HRP anti-mouse polyclonal goat Dako 1:2000, HRP anti-rabbit polyclonal swine Dako 1:2000) diluted in 5 % non-fat milk TBS-T or 1% BSA TBS and subsequently developed with the ECL system (GE Healthcare). Bands were quantified by densitometry using Gel Pro software and were normalized to the loading control.

#### Genetic variant analysis of *ARMCX3*

Genetic variants, loss of function mutations and allelic frequency (see Table S1) of (NRNPH2, ARCMX4, ARMCX1, ACRMCX6, ARMCX2, ZMAT1, ARMCX5) *ARMCX3,* the flanking genes (*NRNPH2, ARCMX4, ARMCX1, ACRMCX6, ARMCX2, ZMAT1, ARMCX5*, per order in human chromosome X); as well as RHOT1, RHOT2 and TRAK2 in control populations were assessed in VARSOME (https://varsome.com/). Involvement in mendelian diseases was determined in Human Mutation Gene Database (HMGD, professional version https://digitalinsights.qiagen.com/products-overview/clinical-insights-portfolio/human-gene-mutation-database/) and in pathogenic structural rearrangements was consulted in ClinVar (in https://genome.ucsc.edu/) (Supplementary Table S1)

#### Statistical Analyses

Statistical Excel (Microsoft), GraphPad Prism (GraphPad), and SPSS Statistics (SPSS) softwares were used to analyze the data. Unpaired Student's t test (with Welch correction when appropriate) or paired Mann-Whitney test were used to test differences between two conditions. Comparison of multiple conditions was performed by one-way ANOVA with Bonferroni post hoc test, for parametric data, or by Kruskal-Wallis test followed by Dunn's correction, for non-parametric data. For the brain width and height measurements, a general estimated equations for repeated measures analysis was performed using the SPSS software. Statistical significance was set at $P < 0.05$, and represented as $*P < 0.05$, $**P < 0.01$, $***P < 0.001$, $****P < 0.0001$. All values in text are given as mean ± s.e.m.
