## Supplementary Figures for "A novel Alex3/Gα_q_ protein complex regulating mitochondrial dynamics, dendritic complexity, and neuronal survival"

**Supplementary Figure 1**

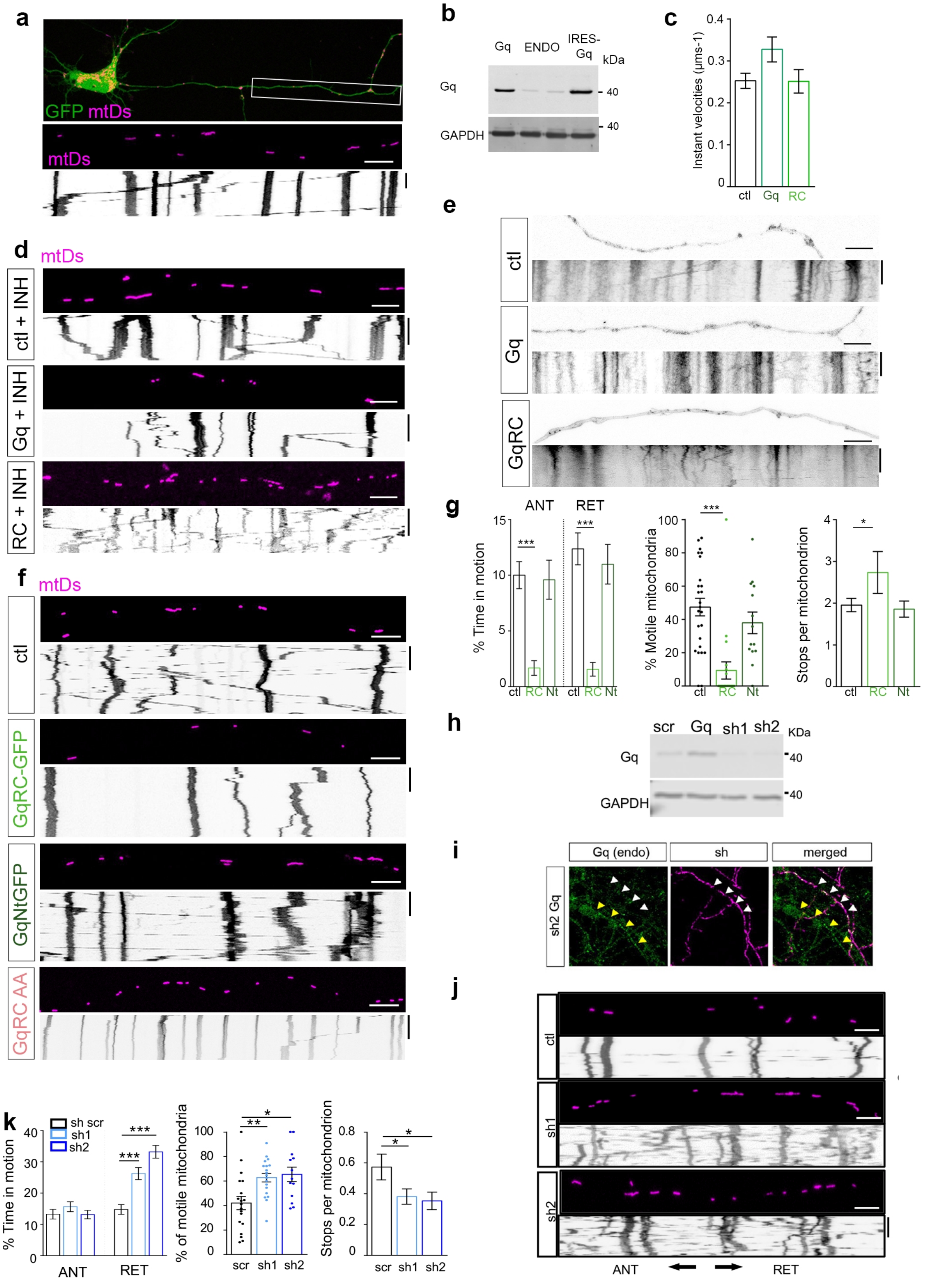

**Supplementary Figure 2**

**a**

| Gene | Peptides C19 WT | Peptides C19 KO+Gq | Peptides C19 NIH 3T3 | Peptides E17 WT | Peptides E17 KO+Gq | Peptides E17 NIH 3T3 |
|---|---|---|---|---|---|---|
| GNAQ/11 | 1 | 5 | 1 | 12 | 12 | 2 |
| WDR36 | | 3 | 11 | | 2 | 24 |
| Cav1 | | | 4 | | | 3 |
| Triobp | | | 4 | | | 3 |
| Plekhh2 | | | | 1 | 2 | 2 |
| Armcx3 | 13 | | | 2 | | 7 |
| Armc10 | 7 | 1 | | 6 | 2 | |

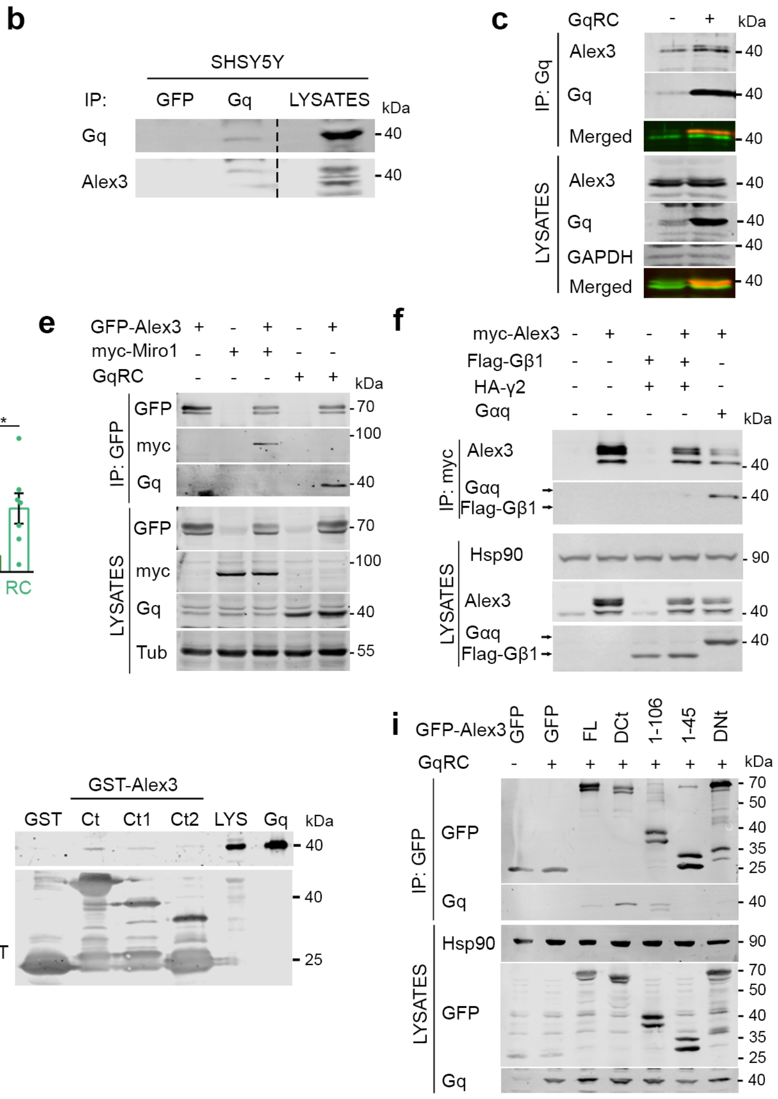

#### Supplementary Figure 3

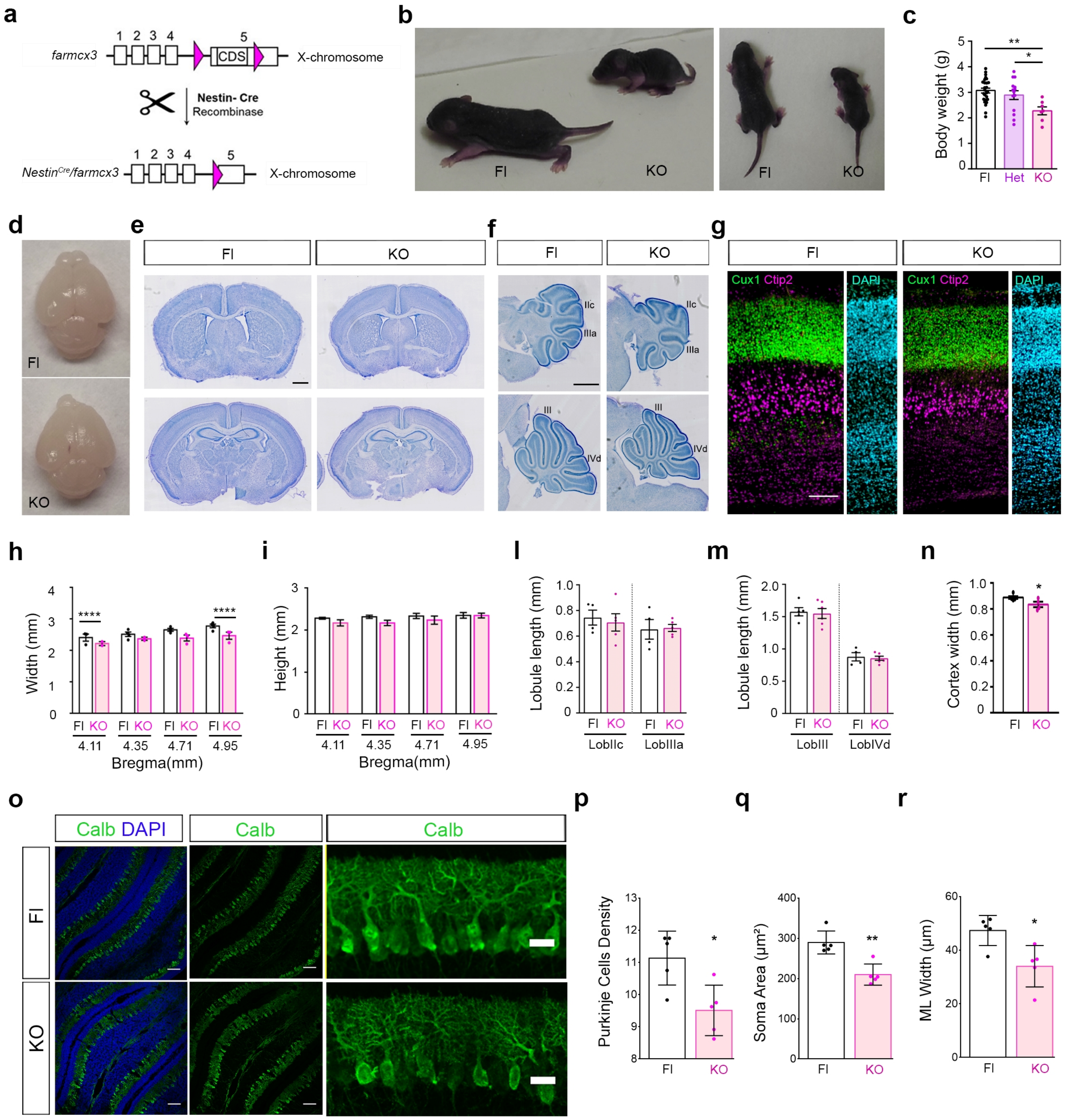

**Supplementary Figure 4**

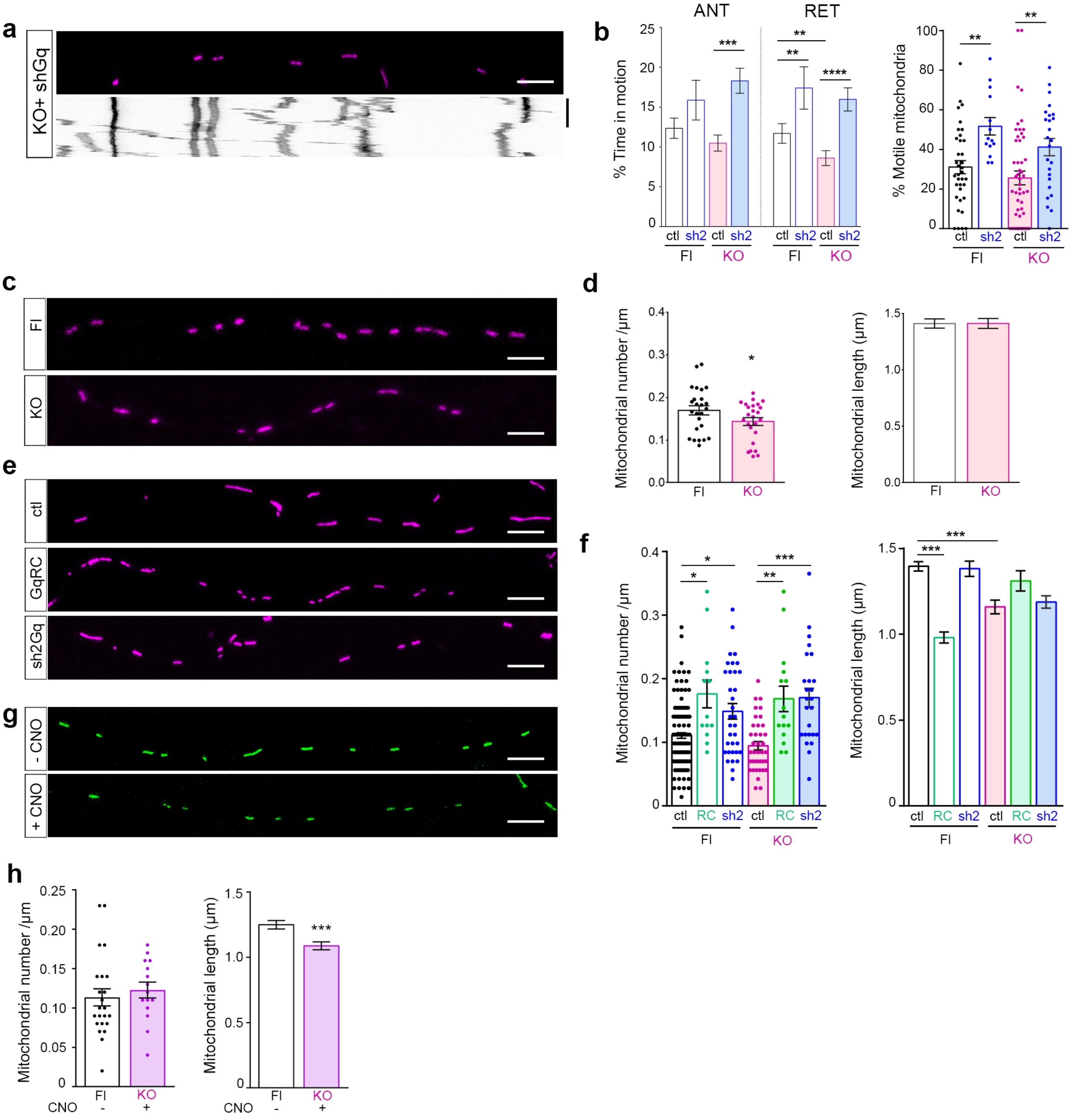

**Supplementary Figure 5**

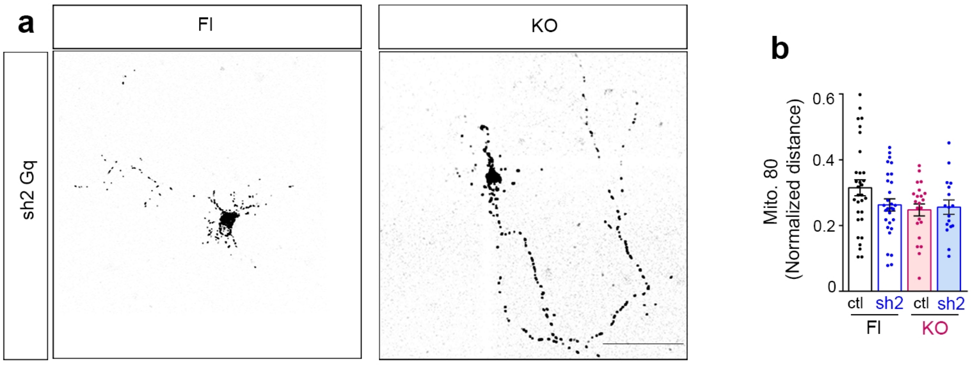

**Supplementary Figure 6**

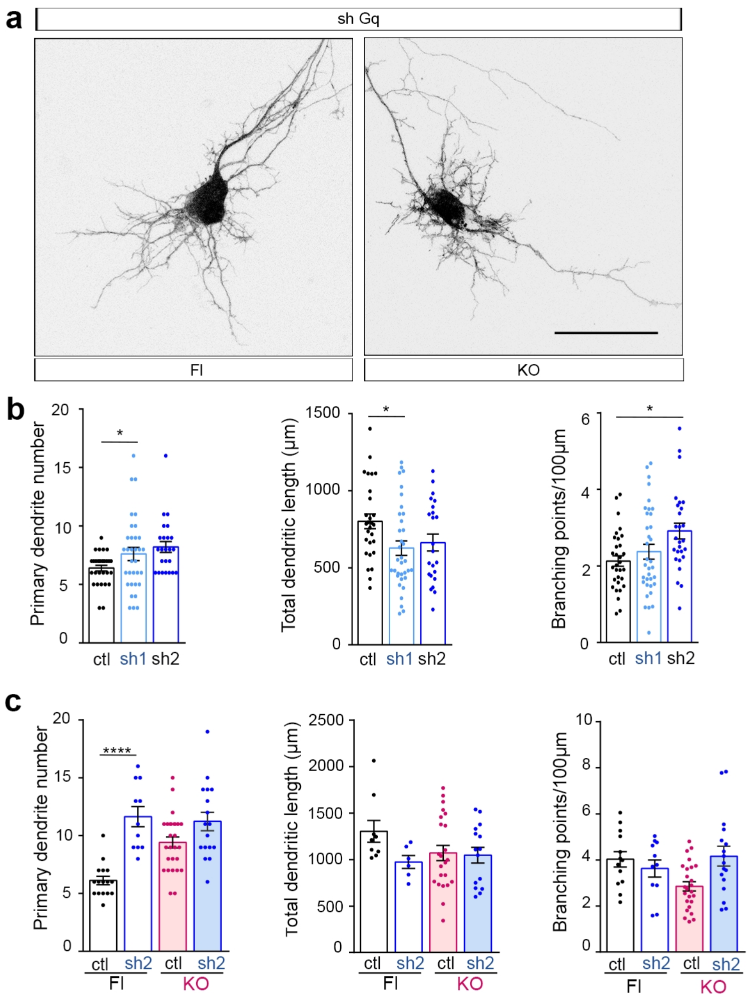

**Supplementary Figure 7**

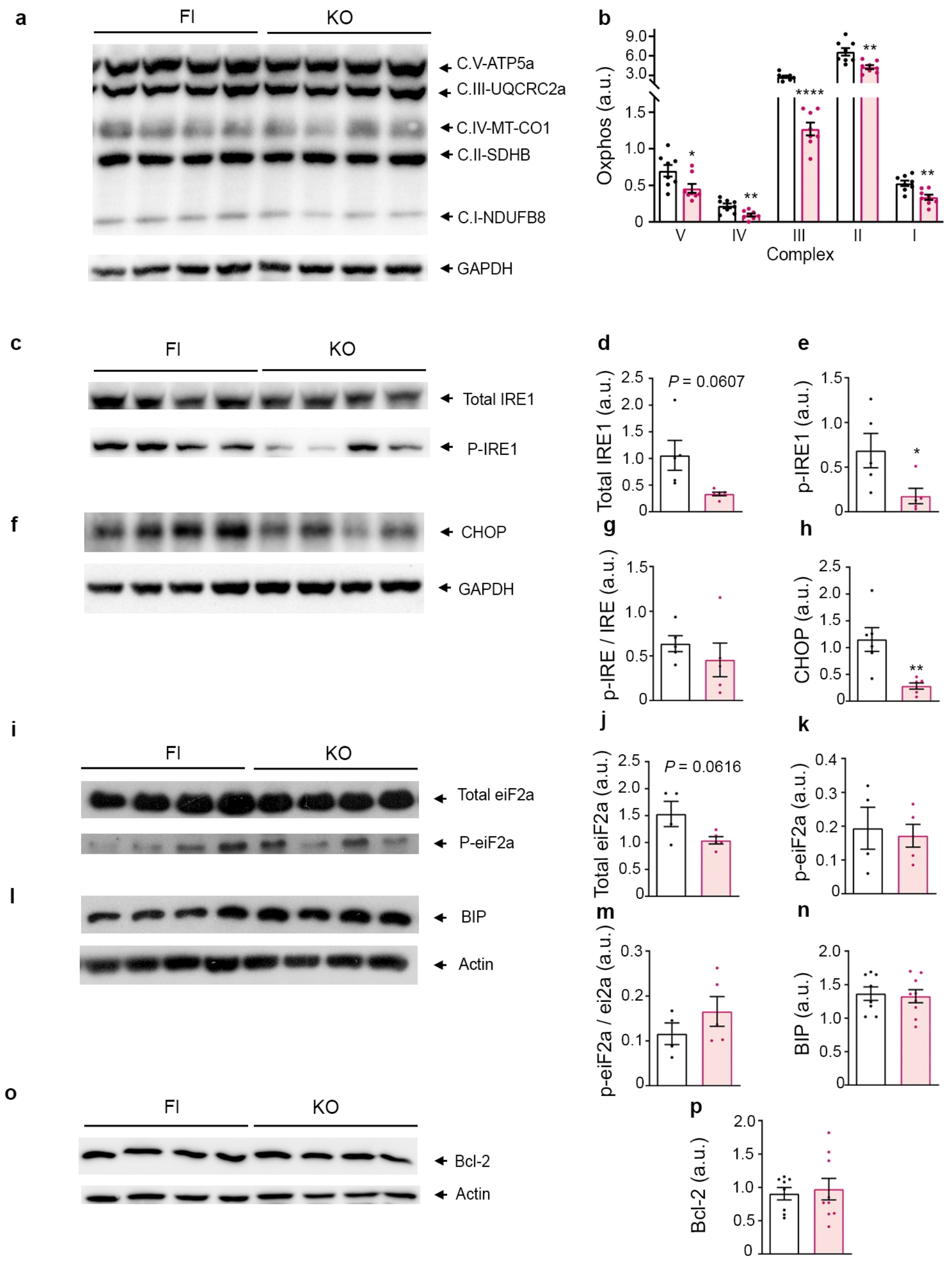

**Table S1 - Genetic variants, loss of function mutations and disease implication of *ARMCX3* and ARCMX cluster genes (by position in chrom. X) versus other genes related to ARCMX3 mitochondrial function ***

| GENE | POSITION | SYNONYMOUS | MISSENSE | LOSS OF FUNCTION (observed/expected) | pLI (probability of Loss-of-function intolerance) | HEMYZYGOSIS (control population) | MENDELIAN DISEASE (causative mutations) # | GWAS (variant association to disease) † |
|---|---|---|---|---|---|---|---|---|
| ***ARMCX4*** | X:100673275-100788446 | 208/282.7 | 559/764.2 | 1/38.7 | 1 | 1/>90.000 | — | — |
| ***ARMCX1*** | X:100805514-100809683 | 67/72 | 107/175 | 3/8.1 | 0.07 | 1/>180.000 | — | — |
| ***ARMXC6*** | X:100870110-100872991 | 47/33.5 | 110/85.6 | 2/5.3 | 0.12 | 16/>180.000 | — | Serum uric acid levels (by proximity with ARMCX3) (5) |
| ***ARMCX3* (Alex3)** | **X:100877787-100882833** | **46/49.6** | **82/140** | **1/5.6** | **0.46** | **None** | — | **Metabolic measurements (uric acid, creatinine), ischemic stroke, body height, sleep-disordered breathing hypoxia (5, 6)** |
| ***ARMCX2*** | X:100910267-100914876 | 115/104.9 | 192/260.8 | 1/8.2 | 0.69 | 3/ >180.000 | — | — |
| ***ZMAT1*** | X:101137262-101187004 | 62/67.6 | 187/204.9 | 8/18.5 | 0 | None | — | Scoliosis (7) |
| ***ARMCX5*** | X:101854096-101859087 | 72/72 | 156/196.9 | 4/10.3 | 0.03 | 1/>180.000 | — | — |
| | | | | | | **HOMOZYGOSIS (control population)** | | |
| ***GNAQ* (Gqα)** | **19:80335191-80646219** | **75/80.1** | **7/80.8** | **2/20.9** | **0.98** | **None** | **Sturge-Weber syndrome (neurocutaneous disorder with brain malformations and intellectual disability (1)** | **Bipolar disorder, body mass index, hepatic protein levels (5, 8, 9)** |
| ***RHOT1* (Miro1)** | 17:30469473-30580393 | 101/124 | 230/362.3 | 9/44.9 | 0.8 | None | Parkinson's disease (2) | Self-reported cognitive performance, nicotine dependence, cobesity (10, 11, 12) |
| ***RHOT2* (Miro2)** | 16:718086-724174 | 267/186.1 | 479/400.3 | 38/34.8 | 0 | None | Autism spectrum disorder, congenital heart disorder (3, 4) | Attention deficit hyperactivity disorder, antisocial behaviour, substance abuse, electrocardiography (13, 14, 15) |
| ***TRAK2*** | 2:202241930-202316302 | 171/178.4 | 458/488.3 | 30/49.4 | 0 | 2/>280.000 | Autism spectrum disorder (3) | Alzheimer disease late-onset, cocaine abuse disorder, carcinoma (16, 17, 18) |

* Data obtained from Varsome (https://varsome.com/)
### Data on involvement of rare mutations in Mendelian disease obtained from Human Gene Mutation Database (professional HGMD)
† Data on assocaition of genetic variants (SNPs) to disease obtained from GWAS Catalog (https://www.ebi.ac.uk/gwas/)

#### SUPPLEMENTARY FIGURE LEGENDS

**Supplementary Fig. 1: G$\alpha_q$ regulates mitochondria motility in axons.**

**a,** Example of 7 DIV hippocampal neuron expressing cytoplasmic GFP and mitochondrial-targeted DsRed (mitoDsRed). An axonal region located 90 to 160 µm from the soma was live imaged for 10 min (white rectangle, middle panel). Kymographs were obtained from resulting movies and mitochondrial motility parameters were determined. **b**, Western blot analysis from HEK293 cell lysates with specific antibodies against G$\alpha_q$ revealed higher levels of protein in cells transfected with plasmids containing G$\alpha_q$ (Gq) or with G$\alpha_q$ bicistronic vectors (IRES-Gq), compared with cells transfected with empty vectors and expressing endogenous protein (ENDO). **c**, Instant velocities of mitochondria from axons (as in Fig 1b) expressing mitoDsRed (ctl), G$\alpha_q$ (Gq) or G$\alpha_q$R183C (GqRC), representing an average speed of motile mitochondria over 0.0083µm/s. n = 402 (ctl), 311 (Gq), and 191 (GqRC) mitochondria velocities averaged. **d**, Representative images and kymographs from axons expressing GFP and mitoDsRed (ctl), GFP and G$\alpha_q$ (Gq), or GFP and G$\alpha_q$R183C (GqRC), in the presence of 10 µM YM-254890 (G$\alpha_q$ inhibitor). For each condition, the first frame of the live-image series appears above the kymographs. **e,** Representative images and kymographs from axons co-expressing the vesicle marker GFP-synaptophysin with mitoDsRed (ctl), G$\alpha_q$ (Gq), or G$\alpha_q$R183C (GqRC). Scale bars, 5 µm. Time bars, 300 s. **f**, Representative images and kymographs from axons co-expressing mitoDsRed with GFP (ctl), the constitutive-active mutant G$\alpha_q$R183C-GFP (GqRC), the GTPase-deficient G$\alpha_q$ (1-124)-GFP (G$\alpha_q$Nt-GFP), or the PLCβ-binding deficient mutant G$\alpha_q$R183C-R156A-T257A-GFP (RCAA). GFP-G$\alpha_q$R183C expression caused a similar effect to that of non GFP-tagged G$\alpha_q$R183. **g**, From axons expressing GFP (ctl), G$\alpha_q$R183C-GFP (GqRC), or G$\alpha_q$ (1-124)-GFP (Nt) along with mitoDsRed, the percentage of time in motion (left), the percentage of motile mitochondria (middle) and the number of stops per mitochondrion (right) were determined and averaged from n = 390 (ctl), 261 (GqRC), and 210 (Nt) mitochondria from n = 25 (ctl), 20 (GqRC), and 14 (Nt) axons. **h,** Lysates (15 ug) of MEF cells expressing GFP scrambled shRNA (scr), G$\alpha_q$ (as control), G$\alpha_q$-specific shRNA (sh1), or a G$\alpha_q$-specific GFP-bound shRNA (sh2) analyzed by western blot with anti-G$\alpha_q$ and anti-GAPDH specific antibodies. **i,** Representative fluorescence micrographs of 7 DIV hippocampal neurons expressing scr shRNA or Sh2 $G\alpha_q$ shRNA. Neurons were stained with anti-$G\alpha_q$ (E17, Santa Cruz) to visualize endogenous $G\alpha_q$. Arrowheads indicate the axon of shRNA-expressing neurons; sh2 $G\alpha_q$ shRNA-expressing neurons barely showed any $G\alpha_q$ signal, whereas non-transfected cells show endogenous signal (yellow arrowheads). Images are maximum intensity z-projections stacks (6 images/stack). **j,** Images and kymographs of axons of hippocampal neurons (7 DIV) expressing mitoDsRed and one of the following: GFP-scrambled shRNA (scr), GFP and sh1, or sh2-GFP $G\alpha_q$ shRNA. **k,** From kymographs as in **j**, the % TIM was averaged for anterograde (ANT) and retrograde (RET) direction. Mitochondria motility was expressed as percentage of total mitochondria moving within axons. n = 136 (scr), 212 (sh1), and 243 (sh2) mitochondria from n = 19 (scr), 19 (sh1), and 13 (sh2) axons from 4 different experiments for each condition.

Results and images are representative of at least 3 independent experiments. Data represent mean ± s.e.m. Statistical analyses: **c,** Kruskal-Wallis with Dunn test statistic = 5.408 $P = 0.0669$; **e,** Kruskal-Wallis with Dunn test statistic %TIM ANT = 41.91 $P < 0.0001$, RET = 50.63 $P < 0.0001$, %MM = 22.82 $P < 0.0001$, Stops = 2.759 $P = 0.2521$**; k,** Kruskal-Wallis with Dunn test statistic %TIM ANT = 3.47 $P = 0.1763$, RET = 47.22 $P < 0.0001$, %MM = 11.77 $P = 0.0028$, Stops = 2.759 $P = 0.2521$.*$P < 0.05$, **$P < 0.01$, ***$P < 0.001$.

**Supplementary Fig. 2: $G\alpha_q$ interacts with Alex3 through its arm domains.**

**a,** Table shows the number of peptides obtained in each condition for the arm-containing proteins Alex3 and Armc10 as well as for $G\alpha_q$ and the previously described mitochondrial partners caveolin 1 (Cav1) and WDR36 after immunoprecipitation and mass-spectrometry analysis. Mitochondria-enriched lysates from MEF WT, MEF $G\alpha_{q/11}$(-/-), MEF $G\alpha_{q/11}$(-/-)+$G\alpha_q$, and NIH3T3 cells were immunoprecipitated using either the C19 or the E17 antibody(10xp150 confluent plates). The IgG fraction was removed, immunoprecipitates were subjected to trypsin digestion, and the resulting peptides were analyzed by mass spectrometry. Data were processed using Scaffold software and proteins containing peptides in the MEF $G\alpha_{q/11}$ (-/-) condition were excluded from further analysis. **b**, Immunoprecipitation of endogenous $G\alpha_q$ from lysates of SHSY5Y cells (700 µg) followed by western blot analysis validated the interaction of Alex3 with G$\alpha_q$ (GFP was used as negative control). **c**, Immunoprecipitation of G$\alpha_q$ from lysates (700 µg) of HEK293 cells expressing or not G$\alpha_q$R183C (GqRC), followed by western blot analysis. Alex3 (green) and G$\alpha_q$ (red) were visualized using an Odyssey infrared system. **d**, Immunoprecipitation of myc-Alex3 from extracts (700 µg) of HEK293 cells co-expressing G$\alpha_q$ or G$\alpha_q$R183C (GqRC) followed by western blot analysis with specific antibodies against G$\alpha_q$, Alex3, and GAPDH (loading control) (left). Levels of co-precipitated G$\alpha_q$ proteins were determined and standardized relative to those of the control condition; data were obtained from at least 4 independent experiments (right). **e**, Immunoprecipitation of GFP-Alex3 from extracts (700 µg) of HEK293 cells co-expressing G$\alpha_q$R183C (GqRC) or myc-Miro1 (used as positive control) followed by western blot analysis with specific antibodies against GFP, G$\alpha_q$, myc, and β-tubulin. **f**, G$\beta_1$ did not immunoprecipitate with GFP-Alex3. Immunoprecipitation of myc-Alex3 from extracts (700 µg) of HEK293 cells co-expressing Flag-G$\beta_1$ and HA-$\gamma_2$ or G$\alpha_q$ followed by western blot analysis with specific antibodies against Alex3, G$\alpha_q$, and Flag. **g**, Schematic diagram of the GFP- and GST-tagged Alex3 constructs utilized in the immunoprecipitation and pull-down experiments. Arm, armadillo-domain; NLS, nuclear localization signal; TM, transmembrane domain; FL, full-length; Nt, N-terminus; Ct, C-terminus; Ct1, C-terminus proximal; Ct2, C-terminus distal; ΔCt, C-terminal-lacking construct; ΔNt C-terminal-lacking construct; 1-106, truncated protein from residues 1 to 106; 1-45, truncated protein from residues 1 to 45. **h**, Pull-down of GST-Alex3 Ct, Ct1, and Ct2 (shown in g) from lysates of HEK293 cells, followed by western blot analysis with specific antibodies against G$\alpha_q$ (endogenous) and GST. 1 µg of purified G$\alpha_q$ and 20 µg of cell lysates were loaded onto Gq and INPUT tracks, respectively. **i**, Immunoprecipitation of the GFP-Alex3 constructs (shown in g) from extracts (700 µg) of HEK293 cells co-expressing G$\alpha_q$R183C, followed by western blot analysis with specific antibodies against GFP, G$\alpha_q$, and Hsp90.

Results are representative of at least 3 independent experiments. Data represent mean ± s.e.m. Statistical analysis: **d,** One-way ANOVA with Bonferroni post hoc test $F(2,15) = 10.72$ $P = 0.0013$..*$P < 0.05$, **$P < 0.01$, ***$P < 0.001$.

**Supplementary Fig. 3: Morphological alterations in the *armcx3* KO mouse line.**

**a**, Generation of *armcx3* floxed/NestinCre mouse line (*Nestin$^{Cre}$/farmcx3*). An *armcx3*-floxed line (farmcx3) was mated to a NestinCre driver line to obtain specific deletion (KO) of *armcx3* in the brain. **b**, P7-8 control (Fl) and KO mice showed noticeable body size differences. **c**, Body weight was significantly lower in KO vs Het and Fl mice. **d**, Representative brains of P7 Fl and KO mice, showing observable brain size differences. **e-f,** Representative images of Nissl-stained brain (**e**) and cerebellum (**f**) sections from P7-8 control and KO mice. **g**, Representative confocal images depicting the cortical distribution of Cux1+ (green, upper layers) and Ctip2+ (magenta, lower layers) cells at P5 in Fl and KO mice; DAPI is in cyan. **h-i,** Quantitative analysis of the maximum width (**h**) and height (**i**) of DAPI-stained brain sections from P5 Fl and KO mice, at different levels respective of Bregma. **l-m**, Quantitative analysis of the maximum length of lobules IIc IIIa, III, and IVa of Nissl-stained cerebellum sections from P7-8 Fl and KO mice. **n**, Quantitative analysis of the cortex width in P5 Fl and KO mice. Note the slight changes in the overall size of the brain, without alterations in lamination. **o**-**r**, Purkinje cells are affected by Alex3 deletion. Representative microphotographs at two different magnifications depicting the distribution of Purkinje (Calbindin+) cells at P8-10 in Fl and KO cerebellum sections; DAPI is in blue, (**o**). Quantification of Purkinje cell number (**p**), Purkinje cell soma area (**q**), and molecular layer width (**r**) in ROI from Fl and KO mice cerebellum sections.

Data represent mean ± s.e.m. Statistical analyses: one-way ANOVA with Bonferroni post hoc test **c**, $F(2, 44) = 6.278$ $P = 0.0040$; n = 7-26 mice per genotype; General Estimated Equations repeated measures with Bonferroni post hoc test: **h,** genotype, Wald Chi square = 32.649 df = 1 $P = 0.0001$; **i**, genotype, Wald Chi square = 3.022 df = 1 $P = 0.082$; unpaired two-tailed Student's t-test, **l,** Lob IIc t = 0.4110 df = 7 $P = 0.6934$; Lob IIIa t = 0.1458 df = 7 $P = 0.8882$; **m**, Lob III t = 0.2692 df = 9 $P = 0.7938$; Lob IVd t = 0.3524 df = 8 $P = 0.7336$; n=4-6 mice per genotype; **n**, t = 2.428 df = 7 $P = 0.0455$ n = 4-5 mice per genotype; **p**, t = 3.172 df = 8 $P = 0.0132$; **q**, t = 4.602 df = 8 $P = 0.0018$; **r**, t = 3.120 df = 8 $P = 0.0142$; n = 5 mice per genotype. $^{*}P < 0.05$, $^{**}P < 0.01$, $^{***}P < 0.001$. Scale bars: **e**, **f**, 0.8 mm **g**, 100 µm, **o,** 20 µm in low-magnification and 80 µm in high-magnification images. Molecular layer (ML).

**Supplementary Fig. 4: The effects of Gq on mitochondrial trafficking depend on Alex3.**

**a,** Representative images of axonal mitochondria in *armcx3* KO hippocampal neurons expressing mitoDsRed and a G$\alpha_q$-specific, GFP expressing shRNA (sh2). **b,** Experiments were done in parallel with samples as in fig 4a, b. From kymographs as in (a) and Fig. 4a, in control (Fl) and *armcx3* KO (KO) axons transfected with the mitoDsRed and either a GFP scrambled shRNA or a G$\alpha_q$-specific GFP shRNA (sh2), we determined the percentage of time in motion (TIM) towards the anterograde (ANT) and retrograde (RET) directions (left) and the percentage of motile mitochondria (MM) (right). n = 431 (Flctl), 126 (Flsh2), 685 (KOctl), and 405 (KOsh2) total mitochondria from n = 36 (Flctl), 13 (Flsh2), 8 (KOctl), and 25 (KOsh2Gq) independent axons. Control samples expressing GFP-scrambled shRNA together with mitoDsRed presented comparable %TIM and %MM values to those of control samples expressing GFP with mitoDsRed control vectors. **c**, Representative images of axonal mitochondria in neurons expressing mitoDsRed from P7 Fl and KO mice. Scale bars, 5 µm. **d**, The number of mitochondria was decreased in KO compared to Fl but no differences were found in mitochondrial length**. e**, Representative images of axonal mitochondria in neurons co-expressing GFP scrambled shRNA, G$\alpha_q$R183C (RC) or G$\alpha_q$-specific GFP shRNA (sh2). **f**, From images as in Fig. 4e, the number of individual mitochondria per µm (left) and their length (right) were determined and averaged. n = 962 (ctl), 416 (RC), 360 (sh2), 256 (KO), 162 (KORC), and 336 (KOsh2) mitochondria from n = 123 (ctl), 13 (RC), 34 (sh2), 38 (KO), 15 (KORC), and 25 (KOsh2) axons. **g,** Representative images of axonal mitochondria in neurons co-expressing mitoGFP and the Gq-specific hM3D DREADD receptor before (CNO-) or 5 min after (CNO+) the addition of 1 µM CNO. **h**, From images as in (**d**), the number of individual mitochondria per µm (left) and their length (right) were determined and averaged. n = 572 (CNO-) and 570 (CNO+) mitochondria from at least 15 different axons each.

Data represent mean ± s.e.m. Statistical analyses: **b**, Kruskal-Wallis with Dunn test statistic %TIM ANT=35.67 $P < 0.0001$, RET = 57.09 $P < 0.0001$, % MM = 19.73 $P = 0.0002$; **d**, mitochondrial number, unpaired two-tailed Student's t-test t =1.834 df =48 $P = 0.0729$ n = 25; **f,** Kruskal-Wallis with Dunn test statistic for mitochondria number=40.46 $P = 0.0001$, mitochondria length = 120.9 $P < 0.0001$; **h,** Mann Whitney test mitochondrial number *P* = 0.2680, mitochondria length *P* < 0.001; mitochondrial length *P* =0.9598 n = 212-243. **P* < 0.05, ***P* < 0.01, ****P* < 0.001.

**Supplementary Fig. 5: Mitochondrial distribution in dendrites depends on Alex3 and Gq.**

**a,** Mitochondrial distribution in control (Fl) or *armcx3* KO (KO) neurons at 7 DIV expressing mtDsRed with or without sh2 shGq (sh2Gq) or scr (ctl). Scale bars, 50 µm. **b,** Length-normalized Mito80 values from Fl or KO neurons at 6-7 DIV expressing mitoDsRed and GFP-scramble shRNA (ctl) with or without GFP-sh2-G$\alpha_q$ shRNA (sh2) from n = 54 (FLctl), 29 (FL+sh2), 22 (KOctl), and 17 (KOsh2) neurons.
Data represent mean ± s.e.m. from 5 different experiments. Statistical analyses: **b,** Two-way ANOVA with Bonferroni correction F(3.95) = 2.1, *P* = 0.0967.

**Supplementary Fig. 6: Alex3 and G$\alpha_q$ regulate dendritic arborization.**

**a,** Confocal micrographs of control (Fl) or *armcx3* KO (KO) neurons at 7 DIV expressing mtDsRed and either GFP-scrambled shRNA (ctl) or GFP-sh2 G$\alpha_q$-shRNA (shGq). **b,** Fl or KO neurons at 7 DIV expressing mtDsRed and GFP-scrambled shRNA (ctl), or GFP with G$\alpha_q$ sh1 shRNA (sh1), or GFP-sh2-G$\alpha_q$ shRNA (sh2). The number of primary dendrites (PD) (left), the total dendritic length (TD) (middle), and the standardized number of branching points (BP) (right) were determined and averaged. n=30 (ctl), 33 (sh1), and 24 (sh2) neurons for PD; n = 28 (ctl), 35 (sh1), and 23 (sh2) neurons for TD; and 31 (ctl), 35 (sh1), and 27 (sh2) neurons for BP. **c,** Experiments were done in parallel with samples as in fig 6f. PD (left), TD (middle), and BP (right) were determined and averaged. n = 16 (Flctl), 11 (Flsh2), 27 (KOctl), and 17 (KOsh2) neurons for PD; n = 9 (Flctl), 6 (Flsh2), 23 (KOctl), 15 (KOsh2) neurons for TD; and n = 13 (Flctl), 11 (Flsh2), 25 (KOctl), 17 (KOsh2) for BP. Control samples expressing GFP-scrambled shRNA together with mitoDsRed presented comparable values as those of control samples expressing GFP with mitoDsRed control vectors. Data represent mean ± s.e.m. from 4 different experiments. Statistical analyses: **b,** Kruskal-Wallis with Dunn test statistic PD = 7.922 *P* = 0.00190, DL = 6.299 *P* = 0.0429, DL = 8.262 *P* = 0.0161. **c,** Kruskal-Wallis with Dunn test statistic PD = 30.00 *P* < 0.001, DL = 3.465 *P* = 0.3253, BP = 10.72 *P* = 0.0134. **P* < 0.05, ***P* < 0.01, ****P* < 0.001.

**Supplementary Fig. 7: Decreased levels of OXPHOS respiratory complexes and altered ER response in *armcx3* KO mice.**

**a**, Representative western blot images of OXPHOS respiratory complexes from brain extracts. **b**, Western blot quantification evidencing reduced levels of all respiratory complexes in KO compared to control (Fl). **c, f, i, l,** Representative western blot images of ER stress related proteins: total and phosphorylated IRE (**c**), CHOP (**f**), total and phosphorylated eiF2α (**i**) and BIP (**l**). **d, e, g, h, j, k, m, n,** Western blot quantification of ER stress related proteins showing decreased levels of pIRE1 (**e**) and CHOP (**h**) and a trend in total IRE (**d**) and total eiF2α (**j**). **o**, Representative western blot image of Bcl-2. **p**, Western blot quantification of Bcl-2 levels showing no differences between KOs and controls. Data represent mean ± s.e.m. Statistical analyses: unpaired two-tailed Student's t-test **b**, CV: t = 2.431 df = 14 $P$ = 0.0291; CIV: t = 3.332 df = 14 $P$ = 0.049; CIII: t = 7.406 df = 14 $P$ < 0.0001; CII: t = 3.583 df = 14 $P$ = 0.003; CI: t = 3.678 df = 14 $P$ = 0.0025; **e**, t = 2.417 df = 8 $P$ = 0.0421; **g**, t = 0.8713 df = 8 $P$ = 0.4090; **j**, t = 2.223 df = 7 $P$ = 0.0616; **k**, t = 0.3321 df = 7 $P$ = 0.7496; **m**, t = 1.151 df = 7 $P$ = 0.2874; **n**, t = 0.2820 df = 15 $P$ = 0.7818; **p**, t = 0.3530 df = 15 $P$ = 0.7290; n = 5-9 mice per genotype; unpaired two-tailed Student's t-test with Welch's correction **d**, t = 2.565 df = 4.112 $P$ = 0.0607; **h**, t = 3.802 df = 5.663 $P$ = 0.01; n = 5-9 mice per genotype. *$P$<0.05, **$P$<0.01, ***$P$<0.001, ****$P$ < 0.0001.
